# Integrated Classification of Cortical Cells and Quantitative Projectomic Mapping Unveil Organizational Principles of Brain-Wide Connectomes at Single Cell Level

**DOI:** 10.1101/2025.05.08.652699

**Authors:** Yun Wang, Hsien-Chi Kuo, Xiuli Kuang, Shenqin Yao, Phil Lesnar, Lydia NG, Yaoyao Li, Laila El-Hifnawi, Natalie Chen, Kasey Zhang, Eric M Li, Yoav Ben-Simon, Songlin Ding, Quanxin Wang, Totte Karlsson, Rachel Delley, Grace Willams, Wei Xiong, Chao Chen, Kai Chen, Zili Huang, Zhi-Feng Yu, Wenjie Xu, Leila Ahmadinia, Sarah Walling-Bell, Julia Andrade, Olga Gliko, Uygar Sümbül, Matt Mallory, Colin Farrel, Ben Sutton, Kelly Jin, Zizhen Yao, Luke Esposito, Susan Sunkin, Lauren Kruse, Qingming Luo, Hui Gong, Anan Li, Jia Qu, Hannah Choi, Stefan Mihalas, Jun Zhuang, Hongkui Zeng, Staci Sorensen

**Author notes:** Equivalent contributions (Yun Wang, Hsien-Chi Kuo, Xiuli Kuang, Shenqin Yao, Phil Lesnar).

## Abstract

Molecularly defined cortical cell types have recently been linked to whole neuronal morphology (WNM), particularly those characterized by whole-brain-wide projections, such as intratelencephalic (IT), extratelencephalic (ET), and corticothalamic (CT) neurons. In contrast, classical morphological classifications (e.g., tufted TPC, small tufted SPC, and stellate SSC) are based primarily on local dendrosomatic and axonal structures, especially apical dendrites. This study bridges these perspectives by establishing a new neuronal taxonomy, analyzing the connectomes of defined cortical cell types, and comparing them with those obtained from bulk anterograde injections. Neurons were sparsely labeled via tamoxifen-inducible Cre lines with GFP reporters, and 1,419 WNM cells were comprehensively reconstructed with Vaa3D-TeraVR from ∼15 areas across six functional regions of molecularly labeled brains imaged with 2p-fMOST. These cells were newly classified by integrating current molecular-WNM and classical morphological perspectives, with sample size augmented by 1,455 publicly available WNM cells reconstructed from the Mouse-Light project and CEBSIT. This effort defined ten combined molecular-WNM-classical morphological cell types: L5ET_TPC, L6CT_NPC, L6b_HPC, and seven IT types—L2/3IT_TPC, L4IT_SSC, L4IT_UPC, L4IT_TPC, L5IT_SPC, L6IT_IPC, and L6IT_car3PC. Clustering, quantitative analyses and random Forest classifier objectively validated these types and revealed their distinct connectomes, along with convergent, topographic, and hierarchical organizations across their projection brain regions. At the single-cell level, multiple organizational principles governing cortico-cortical (C-C) and cortico-subcortical (C-subC) connectomes emerged with unprecedented detail, offering a precise GPS-like tool for *in vivo* recordings and robust datasets for neuronal network modeling. Comparisons with bulk anterograde injection data underscored the limitations of traditional methods in identifying projection targets. Overall, our approach provides significant insights into cortical circuitry and elucidates the complex interplay between neuronal molecular identity, whole morphology, and classical morphological classification.

## Introduction

The mammalian neocortex comprises diverse neuron types that form distinct circuits and serve specialized functional roles. Recent advances in neuroscience have deepened our understanding of molecularly defined excitatory neuronal types in the neocortex, particularly those characterized by whole-brain-wide projections, such as including intratelencephalic (IT), extratelencephalic (ET), and corticothalamic (CT) cells [1–5] (see **Supplementary Table 1** for abbreviations). These major molecularly defined neuron types have been linked to whole-neuron morphology (WNM) of single cells [6]. Recent findings further identified cell types that are consistent across cortical areas, as well as those unique to specific regions, highlighting molecular diversity and subtypes within each category [1, 7, 8]. However, reconciling transcriptomic classifications of cortical excitatory neurons with regional and morphological diversity remains an open challenge. Transcriptomics alone is insufficient to fully define cell types, particularly regarding regional specificity, topography, and variability. Even within a single molecularly defined type, such as IT cells, substantial variations in dendritic and axonal arborization may correspond to distinct functional roles [5–7]. To address this limitation, it is essential to establish a unified classification framework that integrates molecular identity with detailed morphological features using large-scale WNM datasets across multiple brain regions.

Over a century, classical morphological classification of cortical neurons, based on histochemical and Golgi staining techniques, has provided a foundational understanding of cortical circuitry. This classification relying on local dendrosomatic and axonal structures, particularly apical dendrites, classified cortical excitatory neurons into tufted pyramidal cells (TPC), untufted or slender tufted pyramidal cells (UPC), thick tufted pyramidal cells (TTPC), small tufted pyramidal cells (SPC), inverted pyramidal cells (IPC), horizontal pyramidal cells (HPC), and stellate cells (SSC), among others [9–34]. These morphological frameworks have been widely used in neuroscience research to elucidate cortical circuit organization and function. Bridging classical morphological types with molecular-WNM-linked cell types to identify various morphological cell types across large brain areas would be highly valuable for neuroscience research.

Our understanding of cortico-cortical (C-C) and cortico-subcortical (C-subC) connectomes has largely come from mesoscale tract-tracing studies [35–37]. Even with neuron type-specific labeling using Cre-dependent AAV injections in transgenic mouse lines, the labeled populations often include multiple molecularly defined neuron types, and their morphological distinctions are not yet resolved. Moreover, bulk anterograde tracing is limited by contaminations and incomplete coverage, complicating data interpretation. These limitations can be overcome by complete single-cell reconstructions across large-scale functional cortical regions.

In this study, we delineate ten molecular-WNM-classical morphologically combined cell types through comprehensive reconstructions of 1,419 molecularly defined full-morph cells across ∼15 cortical areas spanning six functional regions from 79 molecularly marked brains, supplemented by 1,455 full-structure cells from ML and CEBSIT public sources (ML: https://ml-neuronbrowser.janelia.org/ & CEBSIT: Digital Brain). These ten cell types were classified by integrating molecular and classical morphological categorizations: L2/3IT_TPC, L4IT_SSC, L4IT_UPC, L4IT_TPC, L5IT_SPC, L5ET_TPC, L6IT_IPC, L6CT_NPC, L6IT_car3PC, and L6b_HPC. Each type was subjectively identifiable by its distinct dendrosomatic, local, and long-projecting axonal structures, particularly its apical dendrites, and objectively validated with high accuracy. Notably, dominant rather than specific expression of a transcriptomic marker aligned with morphological types. Systematic quantitative analysis of axonal projections grouped these cell types into four connectomic categories: cortical-dominant, subcortical-dominant, L4-cell originating, and L6-cell originating. Within these groups, laminar distribution and target numbers varied significantly across individual cell types. Distinct patterns also emerged in the highly convergent TH, CP and MO regions, where axonal clouds from various cortical areas densely intertwined and functional modular structures were formed with both specialized and highly integrated hubs. Importantly, single-cell topography revealed distinct spatial organization principles governing cortical and subcortical connectivity. Furthermore, hierarchy scores derived from single WNM data correlated significantly with target number and targeting probability but not with targeting strength, and most cell types exhibited clear feed forward (FF) or feedback (FB) preferences. Finally, WNM single cells offered some distinctive advantages over bulk injections in detecting projection targets. Collectively, this approach bridges neuronal molecular-WNM, and classical morphologies, enhancing our understanding of cortical circuitry, and offers a unique and precise tool for *in vivo* recordings and future large-scale neuronal network modeling.

## Results

### Classification of 10 critical cell types in 15 functional cortical areas

Using 79 mouse brains across 25 transgenic lines, we reconstructed 1,419 full morphological excitatory cells from ∼15 cortical areas spanning six functional regions— prefrontal cortex (PFC), lateral cortex, motor cortex (MO), retrosplenial cortex (RSP), somatosensory cortex (SS), and visual cortex (VIS)—using the Vaa3D-VR system (**Figure 1a, b; Supplementary Tables 2, 3**). Neocortical excitatory cells were selected for complete dendritic and axonal reconstruction by prioritizing clear labeling over targeting specific molecular cell types. In some cases, cells of a particular type were pre-selected based on their distinct somatodendritic and axonal morphologies as long as available in brains labeled with different genetic identities.

**Figure 1.**
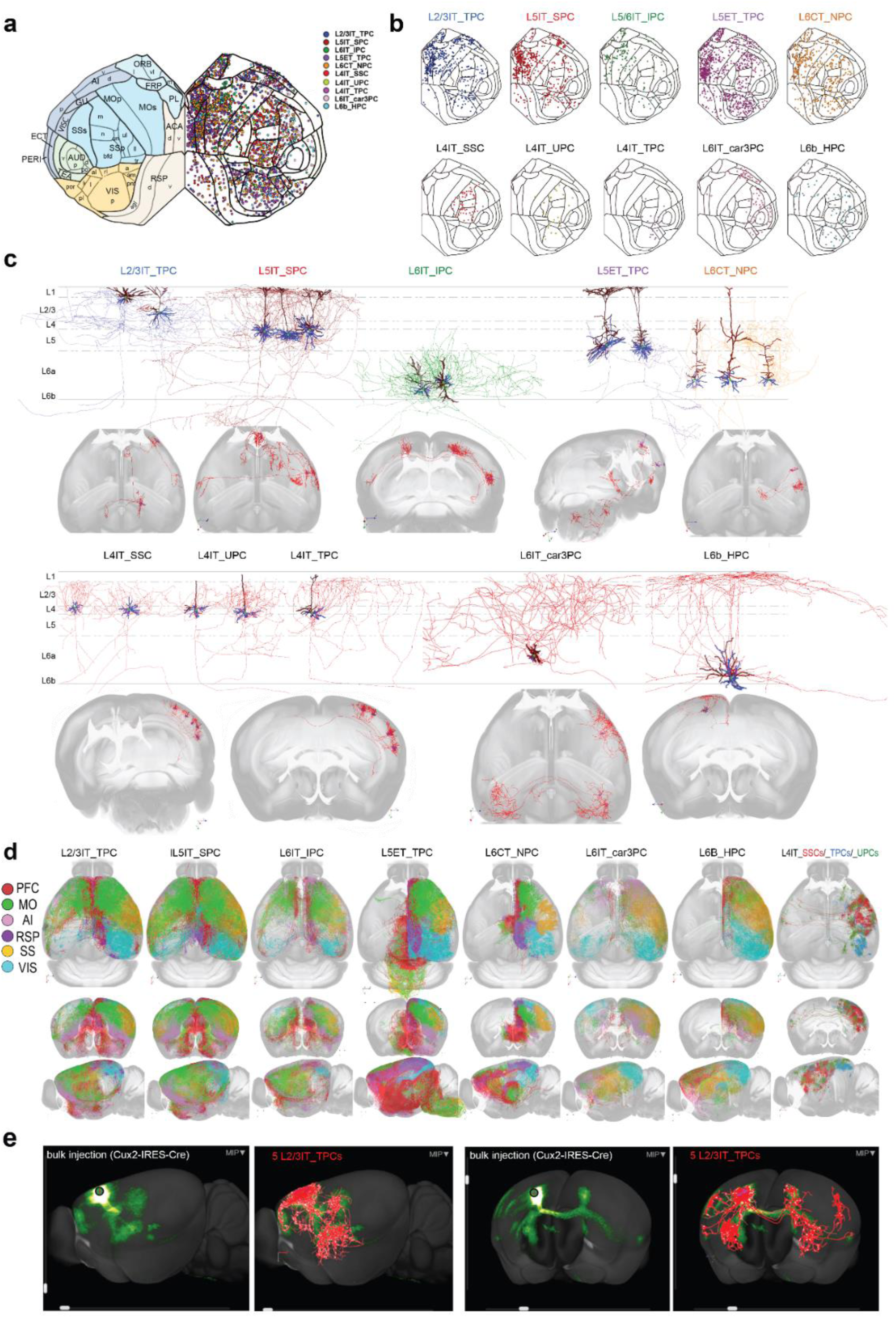
Local and full morphologies of ten cell types across 15 cortical areas and comparison of long-range projections between single WNM cells and anterograde bulk injections. **a.** Soma distribution of all studied cells displayed on a neocortical flatmap of mouse CCFv3, color-coded for individual cell types (all cells flipped to right hemisphere for clarity). Neocortical areas are labeled on the left side with six cortical regions color-coded differently. **b.** Soma distributions of the ten cell types, using the same color coding as in (**a**) on the flatmap. **c.** Local and full morphologies of example cells for each cell type are shown within a schematic diagram of the six-layer cortical structure and in the whole-brain view of 3D mouse CCFv3. Apical dendrites are shown in dark red, basal dendrites in blue. Axons of the five cell types in the upper rows are color-coded as in (**a**), while axons of the five cell types in the lower rows are shown in red. **d.** Full morphologies of ten types of all studied cells, color-coded by six functional regions where somata are located, shown in horizontal, coronal, and sagittal views of 3D mouse CCFv3. As an exception, three L4 cell types are presented together and coded with distinct colors (L4IT_SSC in red, L4IT_TPC in blue and L4IT_UPC in green). **e.** The axonal projections of only five L2/3IT_TPCs in MOs (in red) largely match the labeled projections of an anterograde bulk injection into MOs of Cux2-IRES-Cre specific for the L2/3IT_TPC (**Left panel**: sagittal view; **Right panel**: slightly adjusted coronal view of Experiment 168002073-MOs, https://connectivity.brain-map.org). The soma locations of five L2/3IT_TPCs match the bulk injection site, which is indicated by a red cross within a black circle.

Our previous research linked molecularly defined cell types to WNM, naming them based on whole-brain-wide projections, such as intratelencephalic (IT) and extratelencephalic (ET) cells [5]. This approach allowed us to identify molecular-WNM types through their full morphological features. By examining dendrosomatic and axonal structures, we integrated molecular-WNM classification with classical morphological categories to define ten distinct cell types: L5ET_TPC, L6CT_NPC, L6b_HPC, and seven IT cell types, including L2/3IT_TPC, L4IT_SSC, L4IT_UPC, L4IT_TPC, L5IT_SPC, L6IT_IPC, and L6IT_car3PC (**Figure 1c, d; Table 1**). All seven IT cell types have bilateral (_bi) and ipsilateral (_ips) subtypes, while L5ET_TPCs include two subtypes distinguished by the presence or absence of medullary projections (_MYproj & _NoMYproj). Additionally, 1,455 similarly detailed cells from public databases (ML & CEBSIT) were included in our analysis. Major cell types, including L2/3IT_TPC, L5IT_SPC, L5ET_TPC, and L6CT_NPC, were present across all 15 cortical areas, while others, such as the three L4IT types, L5ET_TPC_MYproj, L6IT_IPC, L6IT_car3PC, and L6b_HPC, appeared selectively in certain cortical regions (**Figure 1b**, **Table 1**). The distributions of a few of these types were found to be largely aligned with genetic labeling results (**Supplementary Figure 1** from AIBS’s public sources: https://lims2.corp.alleninstitute.org; see also *Extended Data* Figure 7 from [3]).

**Table 1.** Key morphological features for identification of cell types and quantitative information of C-C and C-subC connectomes.

|  | L2/3IT_TPCs | L5IT_SPCs | L6IT_IPCs | L5ET_TPCs | L6CT_NPCs | L4cells | L6IT_car3PCs | L6b_HPC |
| --- | --- | --- | --- | --- | --- | --- | --- | --- |
| n = | 563 | 561 | 174 | 811 | 441 | 113 | 103 | 116 |
| typical soma location | L2, L3 | L5 | mainly L6a, also L5 | L5 | mainly L6a, also L5 or L6b | L4 | mainly L6a, also L6b | mainly L6b, also L6a |
| source region selection** | universal | universal | predominantly in PFC, AI and MO | universal | universal | SS, VIS, ADU | rarely in ACA, MO, RSP, VISp | rarely in ACA, RSP |
| classical name | pyramidal cell in L2/3 | small tufted pyramidal cell (SPC) | inverted pyramidal cell (IPC) | thick tufted pyramidal cell (thick TPC) | cortical-thalamus cell (CT cell), narrow pyramidal cell | stellate cell (SSC), untufted pyramidal cell (UPC), tufted pyramidal cell (TPC) | deep layer pyramidal cell | horizontal pyramidal cell (HPC) |
| bilateral ( _bi ) & ipsilateral ( _ips ) subtype, or other subtypes | _bi & _ips | _bi & _ips | _bi & _ips | L5ET-TPC: _MYproj & NoMYproj | n/a | L4IT-SSC: _bi & _ips; L4IT-UPC: _bi & _ips; L4IT-TPC: _ips | _bi & _ips | _ips |
| ratio of _bi_ _ps, or other ratios | 1.9 | 9.2 | 3 | ratio of _NoMYproj:MYproj = 5.2 | n/a | 0.1 (L4IT-SSC); 0.1 (L4IT-UPC); 0.3 (L4IT-TPC) | 0.5 | n/a |
| identical dendritic features | L2IT-TPC: >1 apicals; L3IT-TPC: 1 apical. Both having tufts reaching pial | a small tufted apical w a few oblique dendrites | an inverted apical dendrite downward to WM, can also have oblique, bipolar, horizontal, normally upward orientated apical dendrites | a thick tufted apical with more oblique dendrites; some have an early bifurcated apical dendrite into two apical-like dendrites with a tuft each. | narrow dendritic clusters, untufted or a small tuft apical dendrite reaching L4, but often reaching L1 in VIS. | L4IT-SSC: no apical; L4IT-UPC: 1apical w/o tuft; L4IT-TPC: 1apical with tuft not reaching pial | horizontally oriented dendritic cluster w 1 to 2 bigger apical-like dendrites | horizontally oriented big dendritic cluster with 1 bigger apical-like dendrite |
| identical axonal features (in source region) | axonal cluster branching mainly around their somata in L2/3 and in L5, largely avoiding L4 and L6. | cluster around soma and collaterals up to layer 1 where extending relatively long distance along pial. | axonal cluster of curvy segments selectively in infragranular layers around soma with a few exceptions having axonal collaterals reaching layer 1 | typically a simple cluster around soma; dominantly projecting into multiple subcortical regions with a complexity varied upon soma locating areas | typically a narrow axonal cluster within deep layers 5 and 6, reaching L4; projecting into subcortical areas, dominantly TH. | axonal clusters mainly in L2,3&4, varied size: within a column range (SSC), broader with projections (UPC, TPC) | axonal clusters ramify in all cortical layers in source area extending to multiple cortical areas and projecting to form symmetrical clusters in contralateral hemisphere. | axonal clusters featured with axonal collaterals vertically or obliquely up extending along the pial for a long distance |
| **Axon laminar distributions** |  |  |  |  |  |  |  |  |
| primarily target layers | 2/3, 5, and 1 | 1, 5, and 2/3 | 6a, 5 | 5 | 5 | 2/3, 4 | 2/3, 5, and 6a | 1, 2/3 |
| secondarily target layers | 6a | 6a | 2/3 | 6a, 2/3, and 1 | 6a, 2/3 | 1, 5, and 6a | 1 | 5, 6a, and 6b |
| rarely target layers | 6b | 6b | 1 | 6b | 6b, 1 | 6b | 6b |  |
| **C-C connectomes** |  |  |  |  |  |  |  |  |
| #target | 37 ± 4 | 51 ± 5 | 39 ± 4 | 18 ± 2 | 7 ± 1 | 17 ± 1 | 37 ± 3 | 24 ± 3 |
| TargProb (%) | 21% ± 2% | 26% ± 2% | 32% ± 2% | 22% ± 3% | 27% ± 3% | 20% ± 3% | 32% ± 4% | 32% ± 4% |
| TargStren (Ax/C, μm) | 4892 ± 253 | 4484 ± 270 | 4949 ± 477 | 3689 ± 212 | 4487 ± 269 | 10410 ± 752 | 6665 ± 641 | 7808 ± 948 |
| **C-subC connectomes** |  |  |  |  |  |  |  |  |
| #target | 8 ± 1 | 10 ± 1 | 6 ± 1 | 57 ± 6 | 26 ± 3 | 1 ± 0.2 | 3 ± 1 | 10 ± 4 |
| TargProb (%) | 19% ± 2% | 30% ± 3% | 35% ± 5% | 19% ± 1% | 26% ± 3% | 18% ± 7% | 23% ± 6% | 29% ± 5% |
| TargStren (Ax/C, μm) | 5507 ± 584 | 7743 ± 676 | 4027 ± 721 | 2139 ± 155 | 2552 ± 288 | 3571 ± 995 | 3113 ± 850 | 3953 ± 438 |

High-quality complete structural reconstructions enabled a faithful representation of axonal projectomes from each cortical area. Notably, axonal projections from just a few fully reconstructed cells closely matched those observed in a bulk anterograde AAV injection experiment with minimal contamination when expressing a genetic marker specific to the same cell type (**Figure 1e**).

By visualizing labeled cells in imaged brains and their 3D reconstructions registered in the Allen Mouse Brain Common Coordinate Framework (CCFv3), ten cortical cell types were identified based on distinctive features, including soma location, dendritic (primarily apical), and axonal morphologies (**Figure 1c**, **Table 1**). While individual cell types were typically associated with specific layers, they occasionally appeared in atypical layers.

**L2/3IT_TPCs** were the predominant IT cells in supragranular layers and could be divided into two subgroups based on soma lamination. **L2IT_TPCs** had multiple apical dendrites forming a broader tuft and exhibited predominantly ipsilateral axonal projections. In contrast, **L3IT_TPCs** had a single apical dendrite with a smaller tuft and primarily bilateral projections. Their axons extended across multiple cortical areas in both hemispheres, with limited projections to subcortical regions (see connectome section below).

**L4IT cells** were mainly found in sensory areas (SS and VIS) and comprised three morphological types: **L4IT_SSC, L4IT_UPC, and L4IT_TPC**. L4IT_SSC was predominant in SS but rare in VIS, whereas UPC and TPC were more frequent in VIS and projected to other regions. Most L4IT cells formed an ipsilateral axonal cluster around the soma, though some projected distally to CP or contralaterally, representing bilateral subtypes.

**L5IT_SPCs** were the dominant IT cells in infragranular layers, typically with a few oblique dendrites and a small tuft reaching the pial surface, which was larger in high-functional areas. Their axons formed a cluster around the soma with long-spanning collaterals extending to layer 1. L5IT_SPCs exhibited both ipsilateral and bilateral projections, with a dominance in bilateral projections.

**L5ET_TPCs** had the largest somatodendritic structures among excitatory cortical neurons. Compared to L5IT_SPCs, their apical dendrites were more branched and formed a broader, more complex tuft. They were subclassified based on distal projections to the medulla into

**L5ET_TPC_MYproj** and **L5ET_TPC_NoMYproj** [38]. The latter was found across all studied areas, whereas the former was mainly in MO, SSp-m, and PFC, with lower occurrences in AI and RSP. Among 47 studied L5ET_TPC_MYproj cells, the majority were traced from Fezf2 (28 cells) and Pvalb (13 cells) Cre-lines (**Supplementary Table 3**).

**L6IT_IPCs** were typically characterized by inverted apical-like dendrites and distinct infragranular axonal distributions. Their somata were primarily located in PFC, AI, and MO, though they also appeared in other regions (**Table 1 & Supplementary Figure 1c**). Unusually, L6IT_IPCs shared dominant labeling by the genetic marker *Penk* with another type, L2IT_TPCs(**Table 2**). In VIS and SS, L6IT_IPCs often showed upward oriented or untufted apical dendrites but remained distinguishable by their characteristic infragranular axonal patterns.

**Table 2.**
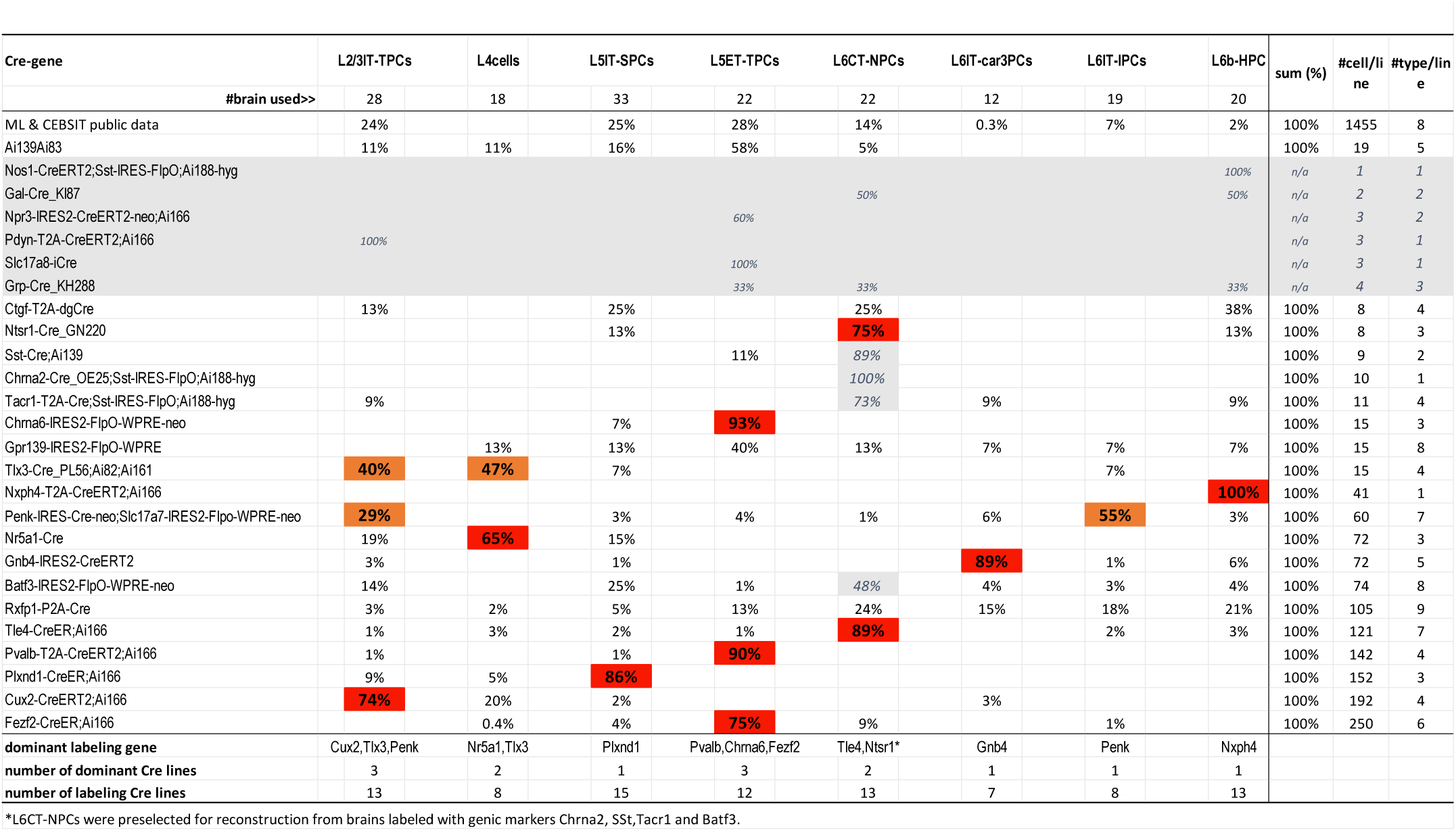
The correlation between Cre-genic markers and full morphorlogical cell types. **Table 2 (Summarized from Supplementary Table 3):**

- Most studied Cre-lines showed dominant labeling of a single cell type (red-shaded, 65–100% probability), though a single Cre-line often labeled multiple cell types, and each cell type was labeled by 7–15 lines. Some Cre-lines labeled specific types with >90% probability. Six Cre-lines (grey-shaded) were excluded from this analysis due to small sample sizes. As Pvalb and Sst are known markers for interneurons (basket and Martinotti cells), only pyramidal cells were selected for reconstruction from these lines to avoid interneuron inclusion.
- Two Cre-lines exhibited dominant labeling of two cell type groups (orange-shaded, 29–55% probability): Penk selectively labeled L2IT_TPC and L6IT_IPC, while Tlx3 labeled L2/3IT_TPC and L4IT types.
- Rxfp1 labels all cell types without strong dominant labeling.
- The labeling probability for L6CT_NPCs appears inflated in four Cre-lines (gray-shaded) due to a preselection procedure based on visually identifiable somatodendritic and axonal structures in brain images.
- Note: 26% of the 1,419 newly reconstructed cells required manual correction of soma location from their originally registered soma locations, primarily for cells near laminar borders or at the cortical bottom, especially in L6b near the white matter or adjacent subcortical structures such as CLA and HPF.

**L6CT_NPCs** had a narrow dendritic cluster with a small tuft or untufted apical dendrite, which remained below layer 3 in most areas but often extended to layer 1 in VIS. Their narrow axonal clusters were largely restricted to deep layers 5 and 6, reaching layer 4 in sensory areas. Cells from sensory regions such as VISp had smaller axonal clusters, whereas those from high-functional areas and MO formed larger, more complex axonal clusters extending to multiple areas.

**L6IT_car3PCs** were primarily located in layer 6a, characterized by horizontally oriented dendritic clusters consisting of a few basal and one or two larger apical-like dendrites, all confined within layer 6. Their axonal clusters spanned all cortical layers within the soma’s location and targeting cortical areas. Uniquely, the bilateral subtype formed largely symmetrical axonal arbors between hemispheres. L6IT_car3PCs exhibited source-region selectivity, largely avoiding PFC, MO, RSP, SSp-bfd, and VISp (**Figure 1b**), a pattern consistent with Car3 gene expression (**Supplementary Figure 1b**).

**L6b_HPCs** resided in the deepest cortical layer and dominantly expressed genes distinct from IT cell types (**Table 2**), closely matching horizontal pyramidal cells (HPCs) in soma location and dendrosomatic morphology[13, 39]. This cell type was featured with long extending axons along pial and ipsilateral projections only, had the biggest dendritic clusters horizontally oriented and restricted within layer 6 or extended partially into layer 5, occasionally further up into supra-granular layers.

Dominant, rather than specific, transcriptomic markers align with morphological types

The relationship between morphological cell types and molecular markers has long been a topic of interest. By refining molecularly defined WNM cell types and reconstructing cells without a preselection procedure, we aimed to establish their correspondence with transcriptomic markers. Among the Cre-lines used to reconstruct 1,419 cells, most exhibited dominant labeling of a single cell type, with a labeling probability—representing the percentage of labeled cells for a specific type within the neocortex for a given Cre-line—exceeding 65%, while a single morphological type was labeled by multiple other Cre-lines with much lower labeling probabilities (**Table 2**). Some Cre-lines showed highly specific labeling, with probabilities exceeding 90%—for example, *Chrna6* selectively labeled L5ET_TPC, *Nxph4* specifically labeled L6b_HPC, and *Gnb4* labeled L6IT_car3PC. Two Cre-lines exhibited dominant labeling to dual cell types: *Penk* dominantly labeled L2IT_TPC together with L6IT_IPC, while *Tlx3* labeled L2/3IT_TPC and L4IT cells. In contrast, *Rxfp1* labeled all cell types without strong dominance for any type.

L6CT_NPCs, characterized by their simple and easily recognizable somatodendritic and axonal structures, were frequently preselected for reconstruction from Cre-lines that primarily labeled other cell types, resulting in an inflated labeling probability for L6CT_NPCs in those lines. For instance, 10 L6CT_NPCs were reconstructed from a brain dominantly labeling Martinetti cells (*ID: 439311-191799, Cre_OE25; Sst-IRES-FlpO; Ai188-hyg*). Similar cases were noted in Batf3, Sst, and Tacr1 lines, which mainly labeled L5IT, Martinotti, and Sst_Chodl cells, respectively. In contrast, when the reconstruction was initially performed without such a preselection procedure from the Tle4 and Ntsr1 Cre-lines—which primarily label L6CT_NPCs—the labeling probability remained dominant at 89% and 75% and at lower than 3% and 13% respectively for other cell types.

Overall, the newly defined molecular-WNM cell types revealed that dominant, rather than specific, transcriptomic markers align with morphological types, underscoring the advantage of our classification approach. This highlights the need for precise morphological refinement to better understand the relationship between genetic identity and cell morphology.

### Validation of the newly defined cell types by objective methods

To objectively validate the WNM cell types identified through human observation of morphological features, we conducted clustering and quantitative analyses on cells across different cortical areas using multiple dendritic and axonal morphological parameters (**Figure 2, Supplementary Table 4a, 4b**). Given the substantial structural variability of cell types across areas, clustering analyses were performed separately for cells in different layers within each cortical area. This approach revealed an overall consistency between the 10 subjectively classified cell types and their objective separation into distinct groups across most of the 15 areas (**Figure 2a**).

**Figure 2.**
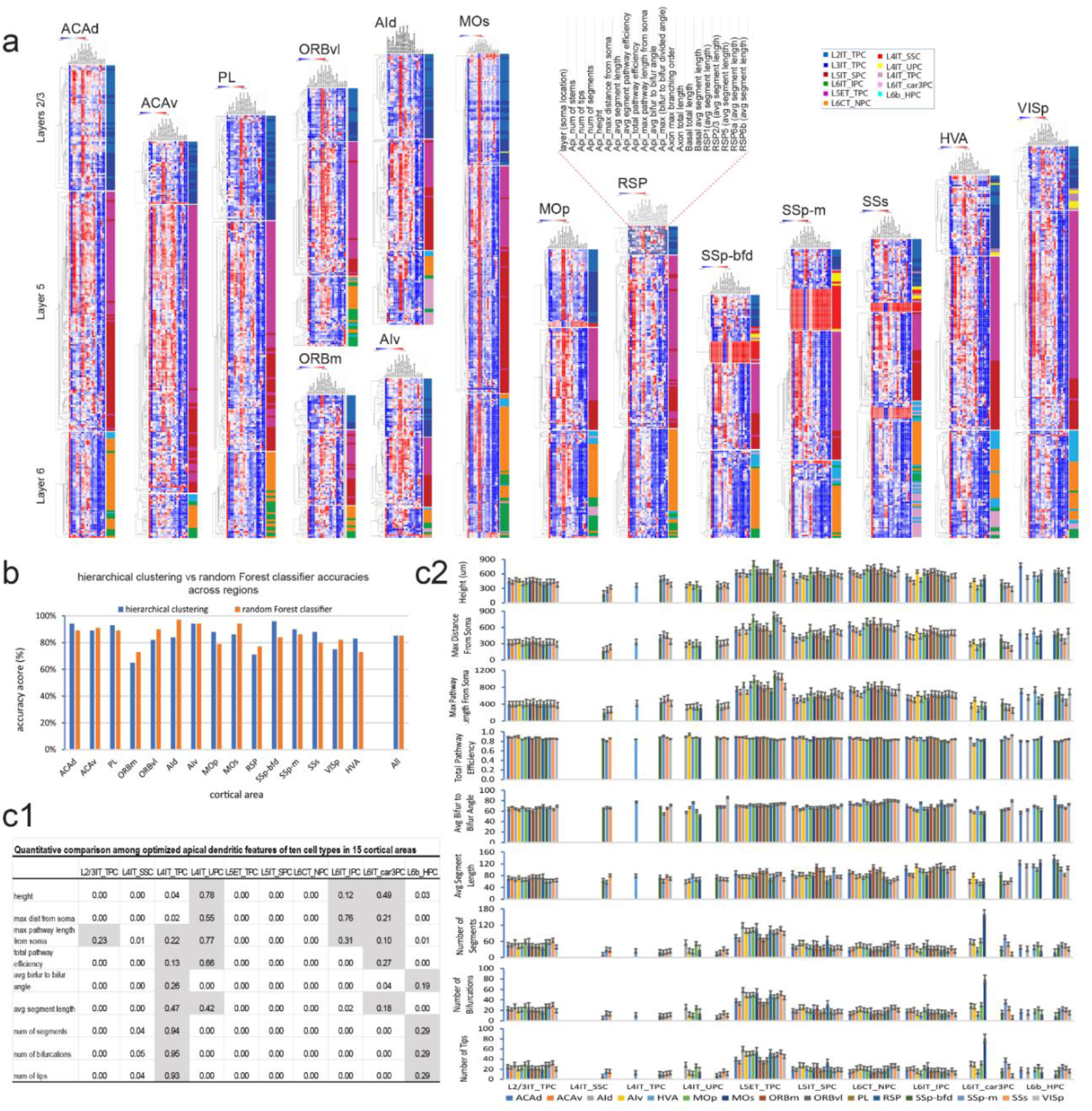
Clustering, quantitative analysis and objective validation of the ten cell types. **a.** Clustering analysis of cell types across 15 cortical areas using One-minus Pearson correlation, based on 21 quantitative features including axonal, apical and basal dendritic metrics, and soma layer location. For each cortical area, clustering was performed separately for three laminar groups: L2–L4, L5, and L6 having L2/3IT_TPCs (further subdivided into L2 and L3 subgroups). Cell types are color-coded on the right of each clustering dendrogram for clearer visualization. **b.** Comparison of hierarchical clustering accuracy and random Forest classifier accuracy across 15 cortical areas and for all pooled cells, showing an overall high degree of agreement between the two methods. **c.** Quantitative analysis of apical dendritic structures across different cell types in 15 cortical areas was conducted using nine selected apical parameters. Statistical comparisons were performed using ANOVA (results shown as P values with those >0.05 shaded, **c1**) and summarized with histogram graphs displaying mean ± SE (**c2**). **Note:** The hierarchical clustering in (**a**) used a broader set of 21 parameters, representing axonal, apical and basal dendritic properties, while the random Forest classifier in (**b**) used a reduced, optimized set of 9 apical dendritic features to maximize classification accuracy and minimize overfitting.

Furthermore, our random Forest classifier, based on optimized morphological features of apical dendrites, successfully predicted cell types across 15 cortical areas with a high average accuracy of 85% (**Figure 2b**). Meanwhile, an ANOVA test showed statistical significance for most parameters of a cell type across 15 areas (P < 0.05, **Figure 2c1, c2**), indicating substantial structural variability likely reflecting the specific functional demands of each brain area. However, for certain cell types—primarily in L4 and L6—the ANOVA results were not significant, indicating that structural differences in their apical dendrites were similar or too subtle to be detected statistically (**Figure 2c1**), as exemplified by the untufted and small tufted dendrites observed in L4 cell types

### Distinctive C-C and C-subC connectomes formed by different cortical types of single WNM cells

Cortical-to-cortical (C-C) and cortical-to-subcortical (C-subC) connectomes are fundamental to understanding brain structure and function. Utilizing complete single WNM cell data, we conducted quantitative analyses encompassing not only neuronal structural characteristics and morphological classification (**Figure 2**, **Supplementary Table 4b**) but also neuronal connectomes (**Figure 3**). These analyses employed key metrics: targeting probability (TargProb), representing the percentage of cells projecting to a specific area, and targeting strength (TargStren), quantifying axonal length within a target area of a given cell type. Target identification criteria were refined to enhance detection of all possible projection targets, particularly weakly targeted areas, which comprised 17% of the total (AxL/C < 1000 µm; see Methods).

**Figure 3.**
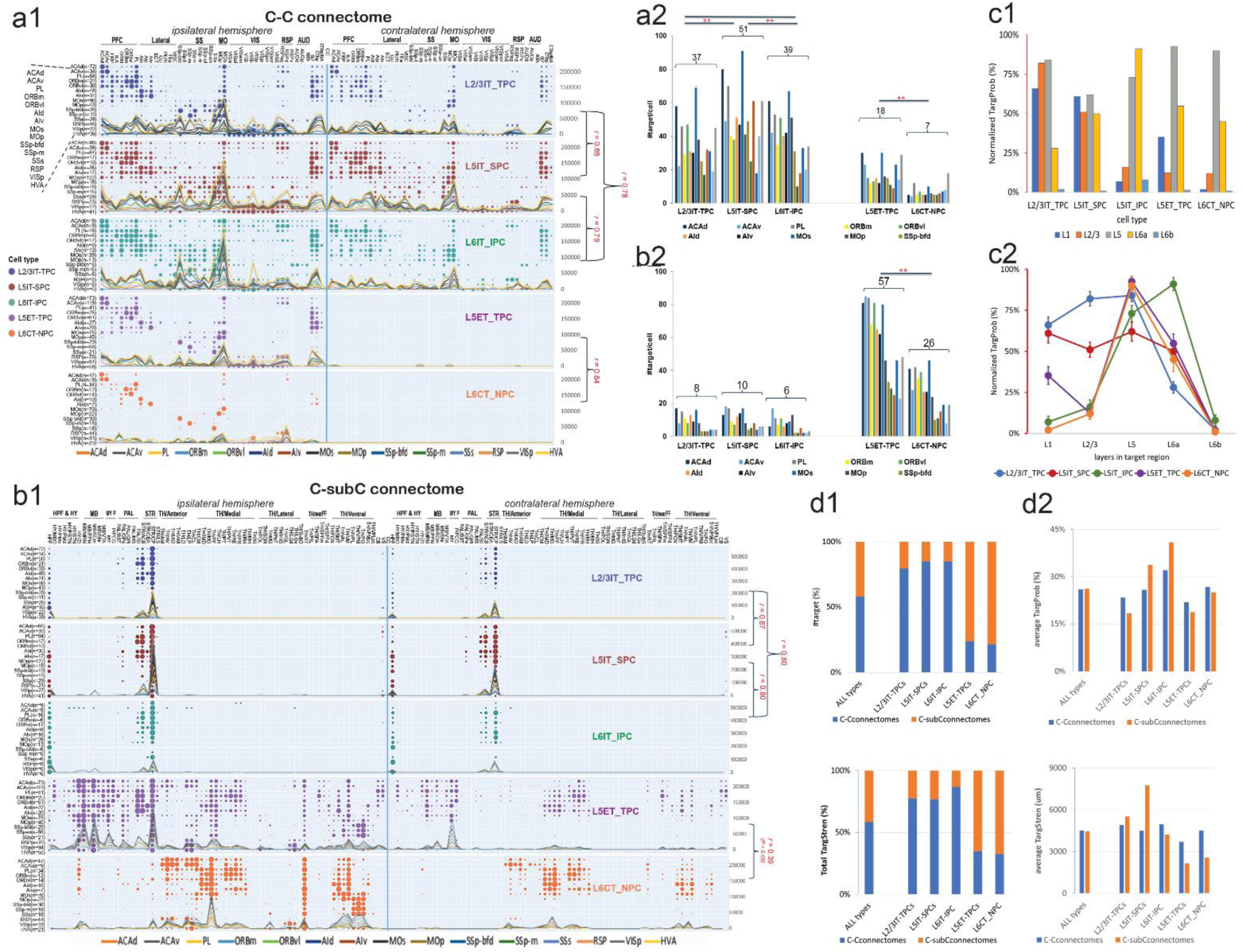
C-C and C-subC connectomes derived from five major cell types: **a.** C-C connectomes of five cortical cell types across 15 cortical areas are shown with TargProb represented as dot plots (%) and TargStren as curvy plots (AxL/C, μm) (**a1**; the same applies to subsequent graph **b1**). The number of targets derived from the C-C connectome by each cell type across different cortical regions (**a2**): The three cortical-dominant targeting types form 2- to 7-folds more targets compared to the two subcortical-dominant types. Among them, L5IT_SPCs produce significantly more targets than L2/3IT_TPCs and L6IT_IPCs. For the subcortical-dominant group, L5ET_TPCs generate significantly more targets than L6CT_NPCs. **b.** C-subC connectome of five cortical cell types across 15 cortical areas (**b1**, utilizing the same cell dataset as in **a1** and **a2**). The number of subcortical targets formed by each cell type (**b2**): The two subcortical-dominant targeting cell types (L5ET_TPC and L6CT_NPC) form 3.7- to 8-folds more targets than the three cortical-dominant types (L2/3IT_TPC, L5IT_SPC, and L6IT_IPC). Among them, L5ET_TPCs generate more than twice the number of targets compared to L6CT_NPCs. In contrast, the three cortical-dominant types consistently form lower and similar numbers of targets (range: 6 – 10). **c.** Distinct laminar distribution patterns of 5 cortical cell types in targeting areas: TargProb (%, mean ± SE) is plotted across five cortical cell types (**c1**) and different cortical layers (**c2**), which is normalized to the maximum laminar TargProb value within each targeted cortical area. Layer 5 is the most intensively targeted, while layer 6b is the least targeted. **d.** Compositions of target numbers and total TargStren in C-C and C-subC connectomes (**d1**) are nearly balanced when pooling all cell types (**All types**), but notable differences emerge at the individual cell types. Similarly, while the average TargProb and TargStren (**d2**) are equal for all pooled types (**All types**), individual cell types show marked variability: higher values of their average TargProb and TargStren are often observed in the non-dominant connectome: for example, L5IT_SPCs, typically exhibiting cortical-dominant targeting, show higher average TargProb and TargStren in the C-subC connectome, whereas L5ET_TPCs and L6CT_NPCs, typically exhibiting subcortical-dominant, present higher average values in the C-C connectome.

All cortical excitatory cell types targeted their soma location areas, forming distinct axonal clusters (∼100% TargProb). Their TargStren patterns correlated with TargProb patterns. Analysis of 2,874 single WNM cells from ∼15 cortical areas revealed four distinct targeting patterns: cortical-dominant, subcortical-dominant, L4-cell originating, and L6-cell originating patterns. The cortical-dominant cell group (L2/3IT_TPCs, L5IT_SPCs, L6IT_IPCs) and the subcortical-dominant group (L5ET_TPCs, L6CT_NPCs) shaped primarily by cell types broadly distributed across all studied cortical areas while the L4-cell and L6-cell groups selectively distributed.

### C-C connectomes

The C-C connectome was primarily shaped by L5IT-SPCs, L2/3IT_TPCs, and L6IT_IPCs, with these three cell types exhibiting similar targeting patterns across both hemispheres (**Figure 3a1, three upper panels**). They projected to multiple cortical areas, with regions such as MOs, AId, OLF, and CTXsp receiving input from nearly all source areas. All three cortical-dominant targeting types projected contralaterally, forming smaller clusters in fewer target areas. Notably, all contralateral targets had corresponding ipsilateral counterparts, resulting in a significant correlation between spatial targeting patterns across hemispheres (r = 0.81, P < 0.01), despite additional targets on the ipsilateral side. However, projections were weaker in the contralateral hemisphere in both TargProb (26% ± 1% ipsilateral vs. 21% ± 1% contralateral, P < 0.01) and TargStren (5235 ± 156 vs. 3348 ± 12 µm, P < 0.01), reflecting asymmetry between hemispheres. On average, these cell types exhibited predominantly bilateral projections, with the _bi/_ips subtype ratio significantly higher for L5IT-SPCs than for L2/3IT_TPCs and L6IT_IPCs (**Table 1**), making L5IT-SPCs the most dominant contralateral cell type. A relatively high proportion of the _ips subtype was found in sensory areas such as VISp and SS.

L5ET_TPCs and L6CT_NPCs, as subcortical-dominant cell types, projected predominantly within the ipsilateral hemisphere, forming axonal clusters near their soma (**Figure 3a1,2, lower panels**). L5ET_TPCs typically exhibited limited collaterals, extending into adjacent areas within the same functional region. However, in higher-order regions such as the PFC and AI, both cell types projected more broadly, frequently targeting MOs, RSP, OLF, and CTXsp.

While sharing similar targeting patterns, notable differences were observed among these targeting cell types, in addition to soma locations, local dendrosomatic and axonal morphologies (**Table 1**). The number of targets per cell varied widely, from 10 (SSp-m) to 91 (MOs), with L5IT-SPCs having the highest target numbers, followed by L6IT-IPCs, while L2/3IT-TPCs had fewer (**Figure 3a2**). In contrast, L5ET-TPCs and L6CT-NPCs exhibited significantly lower target numbers in the C-C connectome (**Figure 3a2**). Further analysis revealed distinct laminar axonal distributions in target areas (**Figure 3c**). L2/3IT_TPCs and L5IT-SPCs preferentially targeted L5 and upper layers, with L5IT-SPCs extending along the pial surface, whereas L6IT_IPCs primarily targeted deep layers. L5ET_TPCs and L6CT_NPCs predominantly projected in their soma-residing layers (L5 and L6).

In the C-C connectome, weak targets showed a correlation between TargProb and TargStren (**Supplementary Figure 2a**). Using a 1000 µm axonal segment length as a threshold[6, 40], 93% of total 2267 targets had axons exceeding this length. Similarly, a TargProb threshold of 2.4% captured 93% of targets, mirroring the pattern observed with the 1000 µm criterion. The remaining 7% of targets, with TargProb below 2.4% and axonal lengths under 1000 µm, represented weak cortical projections.

### C-subC connectomes

The C-subC connectome was primarily shaped by subcortical-dominant cell types, L5ET-TPCs and L6CT_NPCs, which exhibited similar targeting patterns across both hemispheres (**Figure 3b1,2, two lower panels**). These cells projected to multiple subcortical areas, with hippocampal formation (HPF), midbrain (MB), pallidum (PAL), CP, and reticular formation (RT) consistently targeted by cells from most source areas. Targeting patterns were notably uniform in thalamic (TH) regions but weaker in other subcortical areas. This variation arose from L6CT_NPCs in some regions, particularly sensory areas, forming a single axonal cluster largely restricted to TH, whereas those from higher order functional areas and MO generated larger or multiple axonal clusters extending to MB, STR, and PAL. Despite this variation, the correlation between these two cell types remained statistically significant (r = 0.39, P < 0.01), indicating an overall similarity in their targeting patterns.

In addition to disparities in soma locations, local dendrosomatic and axonal morphologies aforementioned (**Table 1**), subcortical-dominant cell types exhibited considerable variation in target numbers, from 8 (VISp) to 85 (ACAv) (**Figure b2**). L5ET-TPCs had significantly more targets, more than double that of L6CT_NPCs. Overall, subcortical-dominant cell types displayed over three times the target numbers of their cortical counterparts (**Figure a2 & b2**). Target numbers generally increased in higher-order functional areas compared to sensory areas, though this trend was less pronounced than in the C-C connectome. In contrast, cortical-dominant cell types (L2/3IT_TPCs, L5IT_SPCs, and L6IT_IPCs) projected to fewer than 10 subcortical areas per cell, primarily targeting STR and HPF, while PAL received much lower TargProb from these cell types in PFC, AI, and MO regions.

Similar to the C-C connectome, a correlation was observed between TargProb and TargStren for weak targets (**Supplementary Figure 2b**). Among 1,749 targets in the C-subC connectome, 69% had axon lengths exceeding 1,000 µm. A TargProb greater than 6% captured the same proportion of targets, mirroring the targeting pattern defined by the axonal length threshold. Thus, 31% of targets had a TargProb below 6% and axonal lengths under 1,000 µm. Compared to the C-C connectome, the total target number was lower (2,267 vs. 1,749), while weak targets were notably more prevalent in the C-subC connectome. Uncommonly, the area RT stood out with a high TargProb but relatively low TargStren, suggesting that subcortical-dominant targeting cell types consistently formed simple axonal branches in RT, maintaining weak yet persistent cortical innervation by all studied cortical areas.

### Overall targeting power balanced between the C-C and C-subC connectomes

Interestingly, the five pooled cell types showed a near balance between the C-C and C-subC connectomes in compositions of both target number (58% vs. 42%) and total TargStren (59% vs. 41%; **Figure 3d1, All types**), as well as in average TargProb and TargStren (26% vs. 26%; 4499 vs. 4431µm, respectively; **Figure 3d2, All types**), indicating a balanced overall targeting power, with both groups contributing to each connectome. Meanwhile, distinct targeting patterns emerged when examining individual cell types (**Figure 3d1, individual cell types**). L2/3IT_TPCs, L5IT_SPCs, and L6IT_IPCs, which dominantly target the cortex, showed higher percentages in the C-C connectome, whereas L5ET_TPCs and L6CT_NPCs, which primarily target the subcortex, exhibited higher percentages in the C-subC connectome. However, this pattern was not directly reflected in the average TargProb and TargStren for individual cell types (**Figure 3d2, individual cell types**), with most types showing trends opposite to their dominant targeting. For example, L5IT_SPCs, primarily cortical-dominant, exhibited higher average TargStren in the C-subC connectome, while L5ET_TPCs and L6CT_NPCs, which predominantly target subcortical regions, displayed higher average TargStren and TargProb in the C-C connectome. This discrepancy may arise from larger axonal clusters or higher TargProb divided by a lower target number, amplifying their average values in these non-dominant areas. For instance, although L5IT_SPCs mainly target cortical areas, their strongest projection was toward CP, a subcortical target. This reversal was offset by the large dominant target number. These findings suggest that specific areas in no-dominant connectome might often be strengthened in the targeting power of a cell type while the overall dominant trend of the cell type was preserved.

### C-C and C-subC connectomes formed by L4 cells

Three distinct types of L4 cells—L4IT_SSC, L4IT_UPC, and L4IT_TPC—were in L4 of sensory areas (e.g., SS and VIS) and were characterized by their small size and predominant _ips subtypes, with _bi subtypes also found in the first two types (**Figure 1c**). In rare cases, L4-like cells were identified in MO and PFC, where layer 4 is not defined in the CCFv3, positioning near the border between layers 3 and 5 [35, 41]. L4IT_SSCs, selectively located in the SS region, often formed compact axonal clusters within a width corresponding to a single cortical column (∼300 µm) [42–45]. This structural feature shaped their projection patterns, primarily targeting layers 2/3 and 4 within the soma’s column, with limited extension to adjacent columns, mostly within SS and MOp (**Figure 4a**). In contrast, L4IT_UPCs exhibited _bi subtypes that shared restricted targeting patterns with L4IT_SSC_bi, while their _ips subtype resembled L4IT_TPC_ips, with broader cortical projections spanning multiple functional regions, including VIS, SS, MO, and RSP.

**Figure 4.**
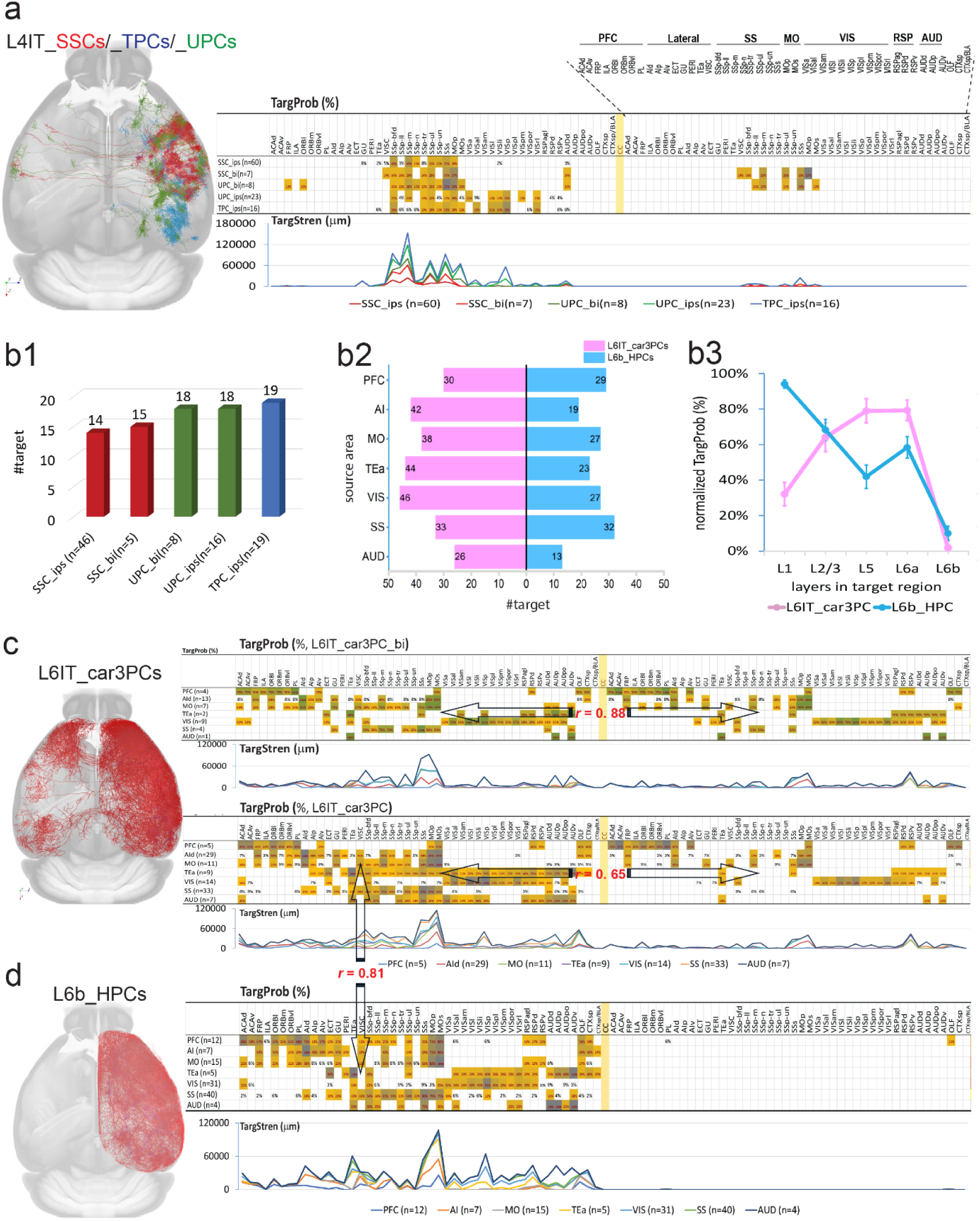
C-C connectomes derived from three L4 cell types and two L6 cell types. **a.** Full morphologies of L4IT_SSCs, L4IT_TPCs, and L4IT_UPCs are visualized in the dorsal view of the CCFv3 template, color-coded in red, blue, and green, respectively (**Left panel**). Notably, each cell type exhibits preferential distributions: L4IT_SSCs in SS areas, L4IT_TPCs mainly in VIS areas, L4IT_UPCs in both SS and VIS regions. Bilateral subtypes are observed at low percentages in L4IT_SSCs and L4IT_UPCs but none in L4IT_TPCs. The C-C connectome of five L4 subtypes is presented with TargProb (%; **Right upper graph**) and TargStren (AxL/C, µm; **Right lower graph**). **b.** Target numbers of five L4 subtypes (**b1**, the same color-code as in **a**), L6IT_car3PCs and L6b_HPCs across seven source areas (**b2**), and distinctive axon laminar projection patterns for the two L6 cell types (**b3**): TargProb (%, mean ± SE) is plotted across cortical layers, which is normalized to the maximum laminar TargProb value within each targeted cortical area. **c.** Full morphologies of L6IT_car3PCs are shown in the dorsal view of the CCFv3 template, color-coded in red (**Left panel**). The C-C connectomes of the L6IT_car3PC_bi subtype and all L6IT_car3PCs (including both _bi and _ips subtypes) are displayed in the same way as in (**a**). Notably, the symmetrical targeting pattern between two hemispheres in L6IT_car3PC_bi connectome results in a high correlation coefficient (r = 0.88), whereas a lower correlation (r = 0.65) reflects a less symmetrical but significant correlation in TargProb of the L6IT_car3PC connectome. **d.** Full morphologies of L6b_HPCs are visualized in the dorsal view of the CCFv3 template, color-coded in red (**Left panel**). The C-C connectome of L6b_HPCs is presented in the same way as in (**a**). A significant correlation coefficient (r = 0.81) is obtained in the ipsilateral targeting patterns between the L6b_HPCs and the L6IT_car3PCs in (**c**).

The average target number of L4 cells in the C-C connectome was 17 ± 3 (range: 14–19), notably lower than that of cortical-dominant targeting types (**Figure 3a2, Figure 4b1**). Their TargProb averaged 20% (range: 14%–31%), also lower than cortical-dominant types. However, their TargStren was 10,410 ± 752 µm per cell—more than twice that of cortical-dominant types (4,773 ± 186 µm per target), suggesting that L4 cells provide the most intensive innervation to their source and nearby areas. All three L4 types projected to the subcortical CP, with TargProb ranging from 6% in L4IT_TPC_ips to higher values in the bilateral subtypes (L4IT_SSC_bi: 43%, L4IT_UPC_bi: 25%).

### C-C and C-subC connectomes formed by other two L6 cell types

Layer 6 contained the most diverse pyramidal cell types. In addition to L6IT_IPCs and L6CT_NPCs, two more types were identified: L6IT_car3PCs and L6b_HPCs (**Figure 1c**, **Table 1**). L6IT_car3PCs were predominantly found in lateral cortical areas (AI, AUD), SS and VIS, while L6b_HPCs were distributed ipsilaterally mainly in SS, VIS, and MO, with rare occurrences in ACA and RSP. To increase sample sizes, cells from TEa and AUD in the lateral cortex were included in the analysis.

L6IT_car3PCs had a significantly higher target number per cell than L6b_HPCs (37 ± 3 vs. 24 ± 3, paired t-test: P < 0.01) (**Figure 4b2**), but their average TargProb (32% ± 4% for both) and TargStren (6665 ± 641 vs. 7808 ± 948) showed no significant differences. Consistent with cortical-dominant targeting cell types, neither exhibited a clear trend of increasing targets from sensory to higher functional areas.

L6IT_car3PC_bi cells (n = 41) displayed a symmetrical targeting pattern between hemispheres (r = 0.88, **Figure 1c**, **Figure 4c**), with no significant differences in target number (ipsilateral: 15 ± 3 vs. contralateral: 11 ± 3), TargProb (54% ± 10% vs. 53% ± 9%), or TargStren (8898 ± 2274 µm vs. 5898 ± 894 µm). However, the predominance of L6IT_car3PC_ips cells (n = 69) resulted in an overall asymmetrical targeting pattern, reflected in significantly higher ipsilateral target numbers (26 ± 2 vs. 11 ± 3, P = 0.01) and TargProb (35% ± 3% vs. 21% ± 6%, P = 0.01), though TargStren differences were not significant (7107 ± 681 µm vs. 5898 ± 894 µm, P = 0.14).

Despite the additional ipsilateral targets, a significant correlation (r = 0.65, P < 0.01) between hemispheric spatial targeting patterns persisted.

L6IT_car3PCs and L6b_HPCs exhibited similar targeting patterns in the ipsilateral hemisphere, with a strong correlation (r = 0.81, **Figure 4c, d**). Both L6 cell types exhibited broad projections across extensive cortical regions. For example, VIS-derived projections spanned all VIS areas and extended into adjacent RSP, AUD, SS, MO, and TEa regions. Notably, the MO region, particularly MOs, was consistently targeted by both cell types from nearly all studied source regions. Significant difference exhibited in their axonal laminar distributions. In layer 1 of target areas, L6b_HPCs exhibited over 2.6 times greater axonal presence than L6IT_car3PCs (84% vs. 34%, **Figure 4b3**), reflecting their characteristic pial-extending axonal collaterals (**Figure 1c**). L6b_HPCs also displayed higher axonal distribution in L6b compared to L6IT_car3PCs (17% vs. 2%).

Subcortical projections were relatively limited. Both cell types, originating from most studied cortical areas, targeted the HPF with an average TargProb of 20% (L6IT_car3PCs) and 28% (L6b_HPCs). Additionally, L6b_HPCs from SS, MO, and ACA projected to the STR, primarily CP, with occasional extensions to ACB and CEA. Some L6b_HPCs, mainly from VISp and MO, targeted different TH areas, while rare instances in AUD, SS, and MO projected to PAL.

Across the four identified connectome patterns, cell types within each group shared common targeting areas, yet their axonal segment distributions varied across cortical layers within a target area. To examine these laminar distribution patterns, we analyzed the TargProb of seven predominant long-projection cell types across individual laminae in five representative target areas (ACAd, AId, MOs, SSp-bfd, VISp) spanning different functional regions of the ipsilateral hemisphere (**Table 1, Supplementary Figure 3**). Each cell type exhibited a distinct laminar distribution pattern across target areas, though the TargProb levels varied depending on their projection sources (**Supplementary Figure 3a, b**; see also **Figure 3c2 & Figure 4b3**). Overall, axonal laminar projection patterns of a cell type showed limited variation among target areas, remaining largely consistent with those in their source areas (**Supplementary Figure 3a, b;** see also **Figure 1c**, **Table 1**).

### Convergent axonal projections in C-C and C-subC connectomes

Convergent projections act as integrative hubs, crucial for complex brain organization and function, and have been extensively studied using anatomical tracing, imaging, electrophysiology, and computational approaches[46–48]. However, they have not been explored from multiple cortical regions using single WNM data. Our analysis of C-C and C-subC connectomes reveals high convergence in CP, TH, and MOs from diverse cell types across all functional cortical regions.

To quantitatively analyze convergent projections, an overlap score was created to measure the overlap between axonal clouds from different source areas, calculated for all 125 pairings among the 15 cortical areas studied (**Figure 5a**). The average overlap score was significantly higher in MOs than in TH and CP (0.79 ± 0.02 vs. 0.69 ± 0.03 and 0.72 ± 0.03, t-test: P < 0.05). Higher-order functional regions generally exhibited greater overlap scores than sensory regions (**Figure 5a1**). Across cortical areas, 76%, 65%, and 74% of pairings had overlap scores greater than 0.5 in TH, CP, and MOs, respectively (**Figure 5a2**), while only 4%, 10%, and 9% showed no overlap, primarily involving sensory regions SS and VIS.

**Figure 5.**
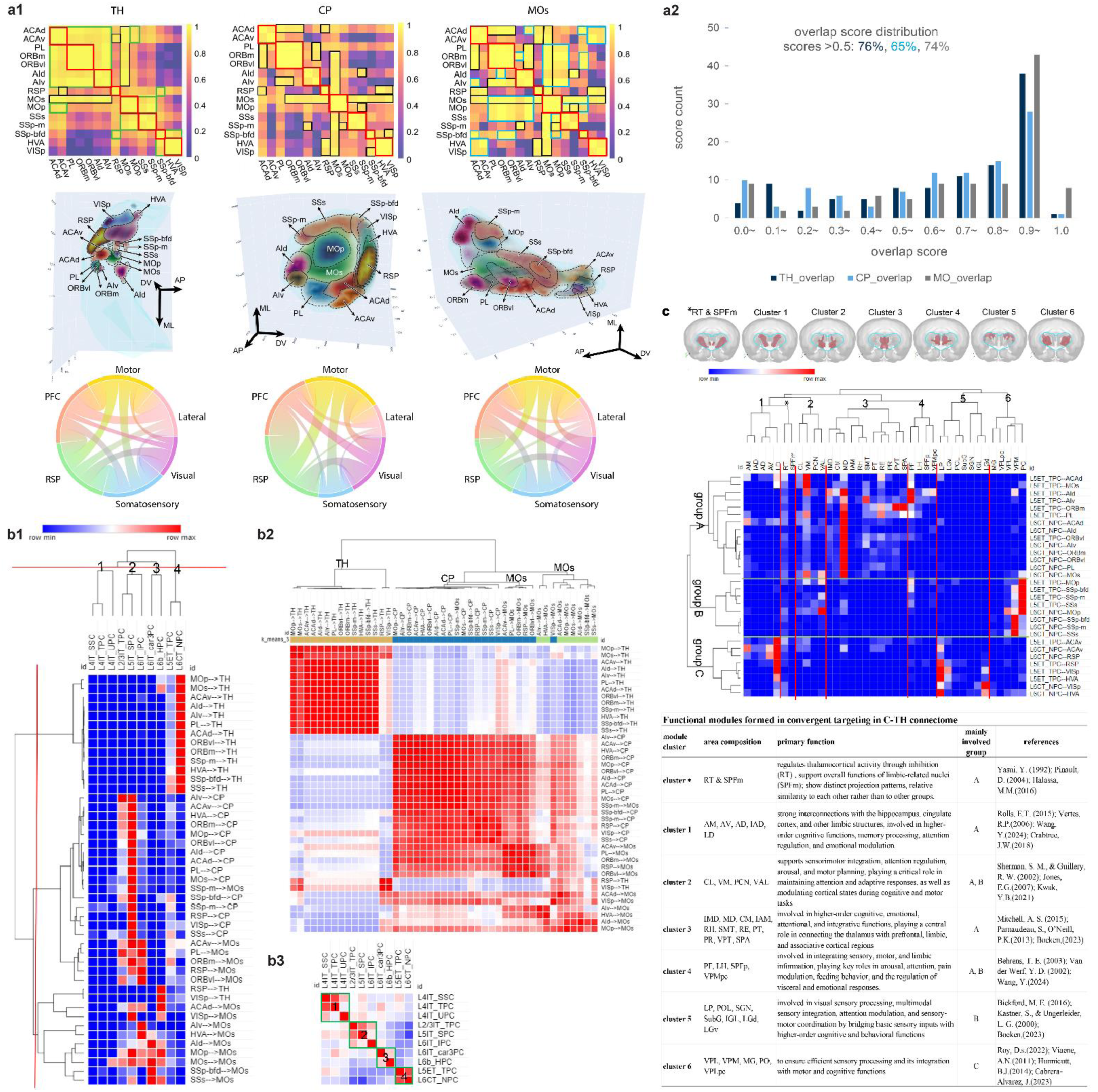
Quantitative analysis of convergent projections in TH, CP and MOs from different cell types across 15 cortical areas. **a. a1-upper panel:** In the overlap score heatmaps, the high-level overlap scores (0.9–1.0) are highlighted among axonal cloud pairs converging in TH, CP, and MOs. Strong overlaps appearing in all three regions are marked with red rectangles, those shared between two regions with black rectangles, TH-specific overlaps in green, and MOs-specific overlaps in light blue. No high-overlap scores were observed exclusively in CP. While CP and MOs display similar heatmap patterns—with MOs showing more high-level overlaps—TH exhibits a distinct pattern, with overlap scores spanning across functional regions such as PFC/AI/MO, MO/SSs, and VIS/SSp-bfd. **a1-middle panel:** Topographic arrangements of axonal clouds from 15 cortical areas are shown in TH, CP, and MOs. To optimally visualize all 15 axonal clouds, central regions were extracted by trimming 75% of the clouds in TH and 40% in CP and MOs. **a1-lower panel:** Overlap score chord diagrams presented for six functional cortical regions all overlap with each other to varying degrees in TH, CP, and MOs. **a2**: Overlap score distributions for TH, CP and MOs are 71%, 68%, and 73%, respectively, exceeding 0.5. **b.** Clustering analysis is derived from the composition of targeting axonal segment lengths in a convergent region from all targeting cell types across 15 individual source areas (**b1**), subsequently similarity matrixes created based on the targets (**b2**) and the cell types (**b3**), respectively resulting in separated clusters reflecting projections in three convergent regions and revealing four cell type groups. **c.** Clustering analysis is derived from axonal segment compositions of two major targeting cell types (L5ET_TPCs and L6CT_NPCs) from 15 source areas across 39 TH subregions, revealing six spatially coherent modular clusters and three major cell groups (A, B, and C). In the top row, the coronal view of the CCFv3 template illustrates the anatomical distribution of these six TH clusters along with RT and SPFm. Subareas within each cluster are spatially adjacent except for RT and SPFm, which exhibit distinct projection patterns. Their relative similarity to each other—rather than to clusters 1–6—may lead the clustering algorithm to group them together. **Inset table**: Functional modules derived from convergent projections in the C-TH connectome with citations of previous studies.

Overlap score heatmaps revealed consistently high overlap scores (0.9–1.0) within the same functional regions, particularly among neighboring areas across TH, CP, and MOs (**Figure 5a1, upper panel**). However, the patterns varied, with TH forming distinct overlap groups, while CP and MOs exhibited similar patterns. In TH, high-overlap groups arose from projections spanning large cortical regions, including PFC/AI/MO, MO/SSs/SSp-m, and SSp-bfd/VISp/HVA. CP shared almost all its high-overlap areas with MOs, but MOs had additional specific high-overlap regions, such as a large group originating from PL/ORB/AI/MO/SSs. Within these highly intertwined axonal structures, projection topographic organization originated from 15 source areas became discernible only after selectively trimming lower-density projections (**Figure 5a1, middle panel**: 75% for TH, 40% for CP and MOs; see **Supplementary Figure 4** for full targeting axonal clouds).

Since cortical areas involved in the same function exhibited high overlap scores and clustered together, we calculated overlap scores across six functional regions (PFC, AI, MO, RSP, VIS, and SS; **Figure 5a1, lower panel**), resulting in 15 possible region pairings for statistical correlation analysis. Notably, CP and MOs showed a strong correlation (r = 0.74, P < 0.05), indicating a similar spatial convergence of axonal clusters, while TH displayed distinct patterns with weaker correlations to CP (r = 0.43) and MOs (r = 0.47). These findings suggest that the spatial organization of axonal clusters in TH, CP, and MOs reflects both functional specialization and a highly integrated convergence of projections.

### Four distinct cell type groups in convergent regions

To better understand targeting dynamics in the three convergent regions, a clustering analysis and a similarity matrix of targets were performed based on the composition of targeting axonal segment lengths from all cell types across 15 source cortical areas (**Figure 5b1, b2**). Cell types projecting to TH clustered together, primarily comprising L5ET_TPCs and L6CT_NPCs from all source areas, with minimal input from L6b_HPCs. Similarly, cell types projecting to CP and MOs tended to cluster together, with limited mixture. L5IT_SPCs from all source areas intensively targeted both CP and MOs, with a stronger preference for CP. L2/3IT_TPCs from most source areas exhibited a similar but weaker pattern. L6IT_IPCs targeted both CP and MOs, with a stronger preference for MOs, particularly from high functional regions and MO, except for VISp and SS areas. L6b_HPCs predominantly targeted MOs, weakly projecting to CP or TH, while L6IT_car3PCs targeted only MOs. The three L4 cell types showed negligible involvement in these convergent regions.

A similarity matrix of cell types revealed four distinct cell type groups (**Figure 5b3**), consistent with previously identified connectome pattern groups at the whole-brain level: the cortical-dominant targeting group (L2/3IT_TPC, L5IT_SPC, L6IT_IPC), the subcortical-dominant targeting group (L5ET_TPC, L6CT_NPC), the L4-cell originating group (L4IT_SSC, L4IT_UPC, L4IT_TPC), and the L6-cell originating group (L6IT_car3PC, L6b_HPC). These findings further validate the classification of WNM cell types in the cortex and highlight distinct targeting preferences and specializations, both at the whole-brain level and within the highly convergent CP, TH, and MOs.

### A broadly integrated modular organization of the C-TH connectome

Looking into the C-TH connectome, 39 of the 44 TH areas reported in this study were targeted either by both major cell types (61%) or exclusively by one (15% by L6CT_NPC and 25% by L5ET_TPC; see **Supplementary Table 5**). L5ET_TPCs formed significantly more targets than L6CT_NPCs (22 ± 2 vs. 19 ± 2, paired t-test, P = 0.05). Both cell types projected to multiple TH areas (average: 21 areas per type from a source area, range: 10-31). Consequently, each target area received a relatively small proportion of axonal clusters, averaging 2.9% from L6CT_NPCs and 1.9% from L5ET_TPCs per source area. The highest percentages were 36.5% of L6CT_NPC axonal segments in VPM from SSp-bfd, and 22.4% of L5ET_TPC axonal segments in LD from ACAv. Notably, RT was targeted by both cell types across all source areas, receiving 3.0% ± 0.3% of axonal segments from L6CT_NPCs and 1.6% ± 0.5% from L5ET_TPCs, with the latter being significantly higher (P = 0.05), indicating a predominant L6CT_NPC innervation in RT. Additionally, MD, PF, VM, VPM, and PO were prominently targeted by one or both cell types from most source areas, with VIS being an exception. VIS was predominantly targeted by both cell types from LD, LP, LGd, and LGv. However, no significant difference was observed in the summed percentages of axonal segment length between the two major cell types from 15 source areas (42% ± 4% for L5ET_TPCs vs. 57% ± 4% for L6CT_NPCs; paired t-test, P > 0.05). These findings suggested that the two major cell types exhibit broadly integrated and overall balanced target strength while L5ET_TPCs innervate more TH sub-areas in the C-TH connectome.

To better understand targeting dynamics across the 39 TH areas, clustering analysis identified six distinct TH clusters targeted by three cell groups: group A (originating from most areas of the PFC, AI, and MOs), group B (from MOp and SS), and group C (from ACAv, RSP, and VIS) (**Figure 5c**). Each group, composed of both L5ET_TPCs and L6CT_NPCs, exhibited distinct targeting patterns with dominant projections to specific TH areas. The six TH clusters, consisting of closely located TH areas (except RT and SPFm), aligned with previously reported functional modules, each associated with specific functional roles (**Figure 5c, inset Table**).

*Cluster 1*, linked mainly to group A, supports higher-order cognitive functions, memory, attention, and emotional regulation through strong limbic connections [49–52]. *Cluster 2*, involving groups A and B, mediates sensorimotor integration, attention, and arousal for adaptive responses [53, 54]. *Cluster 3*, largely associated with group A, integrates higher-order cognitive, emotional, and associative functions [55–57]. *Cluster 4*, connected to groups A and B, processes sensory, motor, and limbic information, contributing to arousal, attention, and visceral regulation [52, 58, 59]. *Cluster 5*, primarily linked to group B, handles visual and multimodal sensory information, integrating it with motor and cognitive functions [55, 60, 61]. *Cluster 6*, associated mainly with group C, ensures efficient sensory processing and its integration with motor functions [62–65]. Notably, all six modular clusters also received weaker projections from other groups, reflecting the highly integrated connectivity of the TH region. These findings align with the overlapping spatial arrangement of axonal clusters in TH (**Figure 5a1, upper panel**), highlighting both functional specialization and broad integration of convergent projections from different cortical regions. Exceptionally, RT and SPFm exhibited distinct projection patterns consistent with their functional differences [66–68], yet were clustered together likely due to their relative projection pattern similarity to each other rather than to clusters 1–6 (**Figure 5c, inset Table**).

In summary, our C-TH connectome analysis reveals a modular organization, with L5ET_TPCs and L6CT_NPCs projecting widely but with balanced strength. Six distinct thalamic modules show strong intra-modular and weaker inter-modular connectivity, highlighting functional specialization and global network integration in C-TH pathways.

### Topographic organization in C-C and C-subC connectomes at the single-cell level

Topographic organization refers to the spatial arrangement of neuronal structures and their functional connectivity within the brain. At the level of single WNM cells, this organization is fundamental to neural circuit function, reflecting signal transmission and network integration. Recent studies have explored the topographic organization of single neurons in select cortical areas, including MO and PFC[40],[6, 69, 70]. To reveal the principles of the topographic organization in large-scale cortical networks, it is essential to examine their broader applicability.

To comprehensively understand the topographic organization of single cells from 15 source areas spanning six functional regions, we analyzed their major targets in both ipsilateral and contralateral hemispheres of PFC, AI, and MO functional regions. Specifically, we correlated the spatial arrangement of somata in a source area with the centroids of their corresponding axonal clouds in target areas (**Figure 6a, b, Supplementary Figures 5, 6**). The orientation of these arrangements was quantified using Spearman’s rank correlation coefficient (**ρ**), measuring the consistency of spatial order (see Methods). Additionally, projection range and percentage were examined, represented by arrow length and thickness in **Figure 6a**.

**Figure 6.**
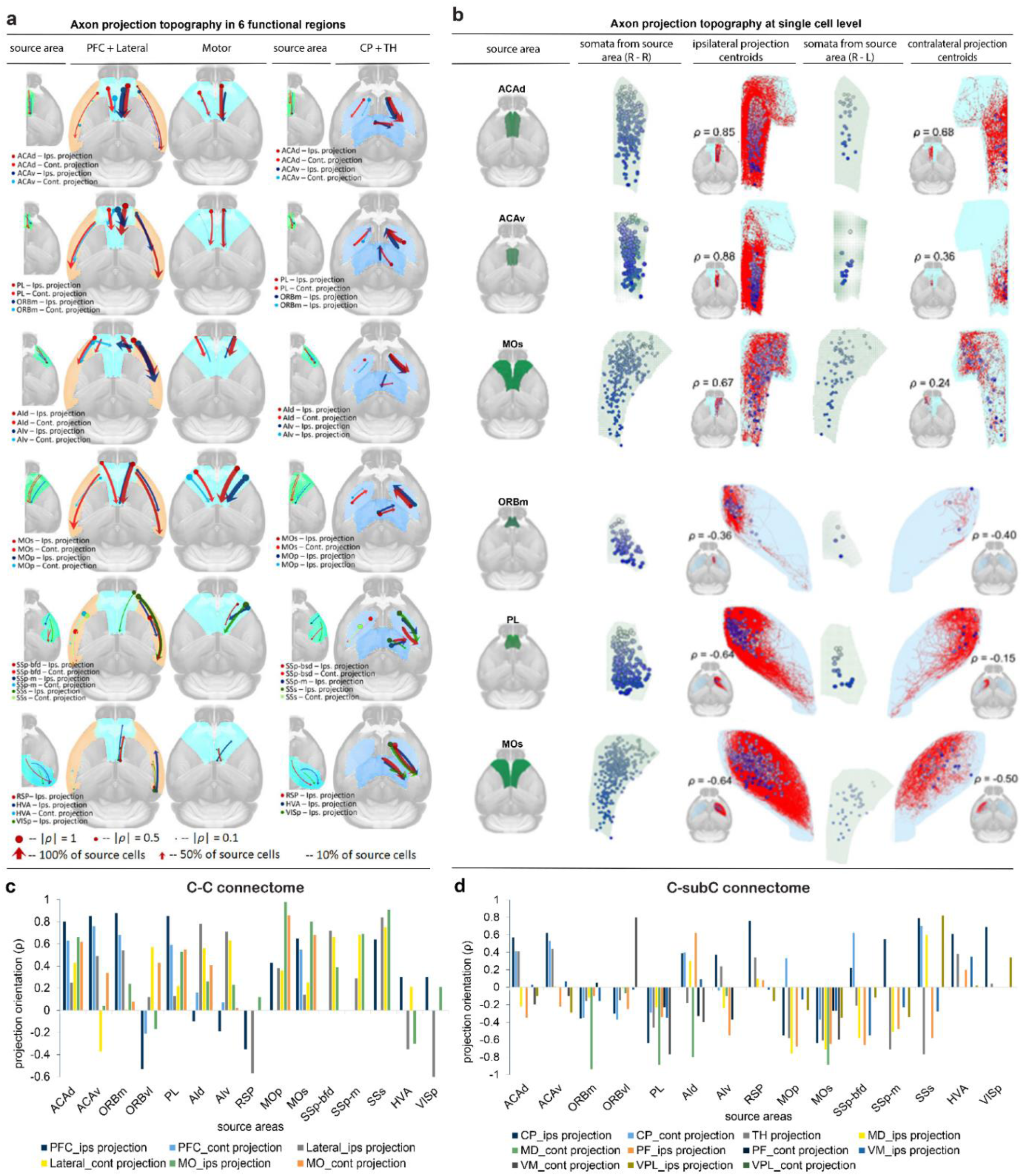
Topographic organization of C-C and C-subC connectomes at the single-cell level. **a.** Axon projection topography was analyzed from 15 source areas to PFC, lateral cortex, and MO in the C-C connectome, and to CP and TH in the C-subC connectome. Spearman’s rank correlation coefficient (**ρ**) quantified the spatial consistency between soma locations and corresponding axonal centroids, with dot size indicating correlation strength. Arrows depict alignment orientation, and their thickness represents the proportion of projecting cells from each source region. **b.** Axon projection topography at the single-cell level is illustrated using three representative examples each for C-C (ACAd, ACAv, MOs to PFC and lateral cortex) and C-subC (ORBm, PL, MOs to CP) connectomes. In these examples, the spatial arrangement of somata in the source region is matched to the corresponding projection centroids in both ipsilateral and contralateral target regions using gradient color density, effectively visualizing the status of their topographic alignment. **c. ρ** value plots for the 15 source areas in the C-C connectome show predominantly positive values with different colors coded for ipsilateral and contralateral projections to PFC, Lateral and MO regions. **d. ρ** value plots for the 15 source areas in the C-subC connectome show predominantly negative values with different colors coded for ipsilateral and contralateral projections to CP, TH, and four sub-areas within TH (MD, PF, VM and VPL).

In C-C targeting regions, axonal centroids largely followed the soma arrangement in the source region (85%, **ρ**: 0.01 to 0.98), with some exceptions (15%, **ρ**: -0.1 to -0.63) (**Figure 6c, Supplementary Figure 5**). In over 50% of source areas, all 6 projection centroids aligned with soma locations, including MO, SS, ACAd, ORBm, and PL. In others, such as ACAv, ORBvl, AI, RSP, and VIS, most projections followed the soma arrangement, but 1-3 projection centroids showed deviations. In contrast, in frequently targeted C-subC regions (e.g., CP and TH sub-areas MD, PF, VM, and VPL), projection centroids often exhibited opposite or divergent orientations (64%, **ρ**: -0.01 to -0.94), with only a minority (36%, **ρ**: 0.01 to 0.79) maintaining the source soma arrangement (**Figure 6d, Supplementary Figure 6**). All projections from PL and MOs were arranged oppositely to their soma locations, whereas those from VISp and HVA aligned with the soma direction.

These findings indicated that while cortical projections generally preserve the spatial arrangement of their source somata, subcortical projections frequently exhibit reversed or divergent orientations.

### Hierarchy organization of brain areas at the single-cell level

The brain’s neuronal networks exhibit a hierarchical organization, with lower-level regions processing sensory or motor information and relaying it to higher-level regions responsible for complex integrative functions, which in turn provide feedback to refine lower-level processes [71] [72–74] [35]. To assess this hierarchy at the single-cell level, we calculated hierarchy scores from WNM data, based on the assumption that hierarchical organization is reflected in the targeting patterns of C-C and C-TH connectomes. Higher-order areas were expected to receive projections more frequently, supported by observations of more complex dendritic structures and broader axonal cloud distributions in these regions [9, 75, 76].

To avoid targeting region size bias, quantitative measurements of TargProb, TargStren, and target number were normalized by axonal cluster size, which enabled a concise calculation of hierarchy scores, producing a distribution curve for target areas (**Figure 7a**; see Methods). In the C-C connectome, higher scores were found in high-level functional regions like PFC and AI, while sensory areas ranked lower. Similarly, in the C-TH connectome, sensory-related target areas in module clusters 5 and 6 (**Figure 5c**) ranked lower, whereas areas involved in higher-order functions ranked higher. Hierarchy score distributions in both connectomes correlated significantly with target number and TargProb but not with TargStren (**Figure 7b**). Notably, some areas deviated from this trend, such as SSp-ul and VISam in the C-C connectome and LGv, LP, PO, and VPL in the C-TH connectome. Despite their primary roles in sensory processing, these areas exhibited higher hierarchy scores, suggesting they receive more frequent projections from source cortical areas than their neighboring regions within the same functional modules.

**Figure 7.**
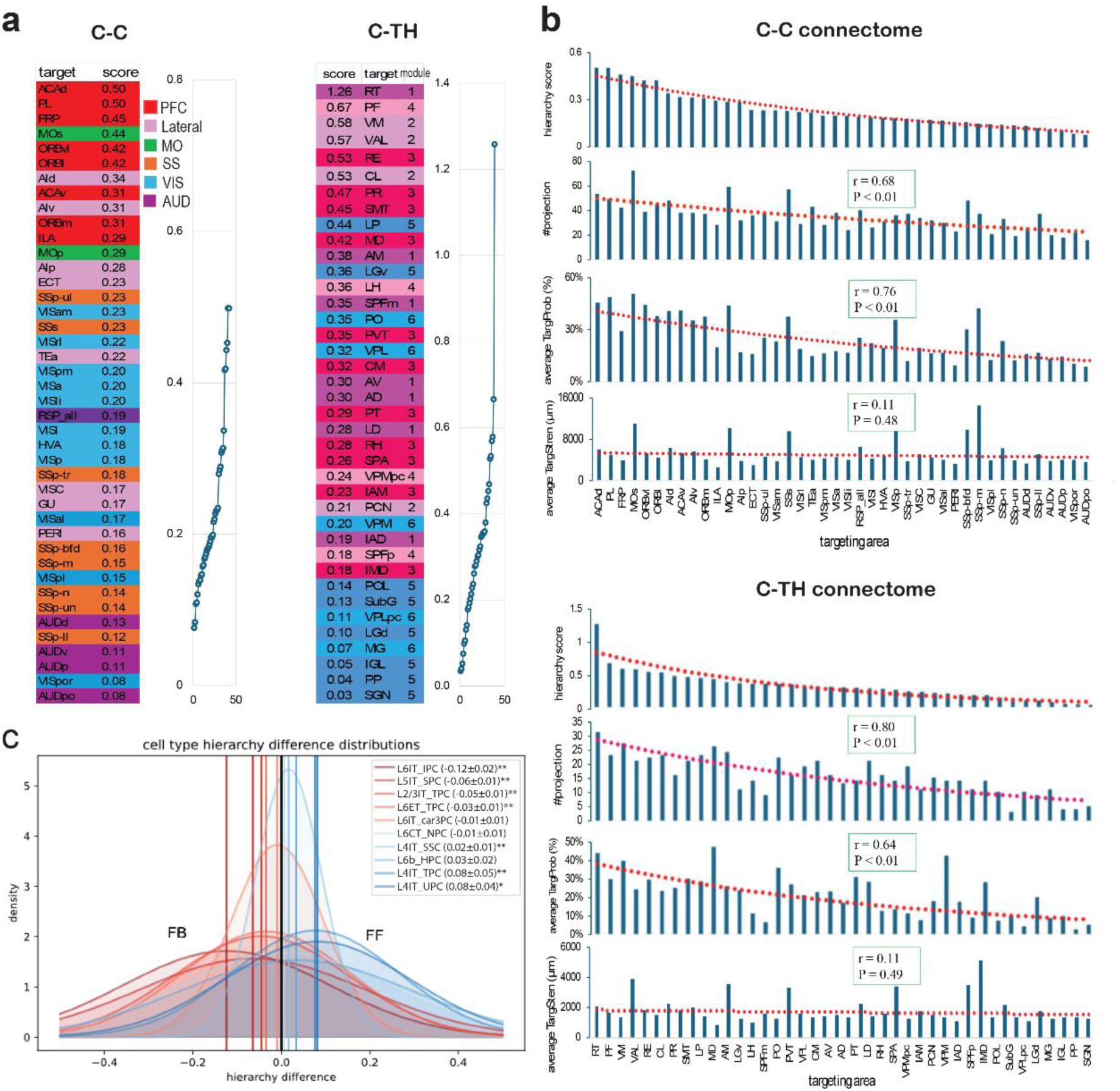
Hierarchical organization of C-C and C-TH connectomes derived from WNM cells across 15 cortical areas. **a.** Hierarchy scores and corresponding curves for different target areas in C-C and C-TH connectomes. The hierarchy score quantifies a brain area’s position within the connectome based on how frequently it is targeted by diverse cell types across 15 cortical areas. In the C-C connectome, higher scores are associated with higher-order functional regions, while sensory areas occupy the lower end of the curve. A similar pattern is observed in the C-TH connectome: areas in module clusters 5 and 6, associated with sensory processing (see **Figure 5c**), show lower scores, whereas areas involved in cognitive, emotional, attentional, and integrative functions rank higher. **b.** Hierarchy scores are significantly correlated with the target number and average TargProb, but not with TargStren, in both C-C and C-TH connectomes. **c.** Hierarchical preferences across cell types. Gaussian fits of the distribution of cell type hierarchy differences reveal FF-leaning types in red and FB-leaning types in blue, with color intensity reflecting the means of the distributions. Vertical lines mark the mean of each fit. **Inset** shows mean ± SE; *P < 0.05, **P < 0.01 (P-values from Wilcoxon tests against variance-matched Gaussians centered at zero).

Our previous bulk anterograde AAV tracing study investigated the hierarchical organization of the C-C, C-TH and TH-C connectomes by modeling connection patterns as feedforward (FF) or feedback (FB), generating testable predictions of hierarchical positions for individual cortical areas and network modules [35]. To classify the FF or FB tendencies of projections from each cell type, we analyzed hierarchy differences in their projections based on a brain area hierarchy constructed from C-C, C-TH and TH-C connections and their layer-specific projection patterns, proposed in [35](see Methods). The distribution of the hierarchy differences for each cell type was tested against a normal distribution centered at zero, revealing significant deviations (P < 0.05, BH-corrected) in 7 of the ten cell types, indicating a clear FF or FB bias with a magnitude reflecting the strength of preference (**Figure 7c**). Notably, L4 cells exhibited the strongest FF tendency, consistent with their axonal projections to L4 and supragranular layers and their somata in sensory regions (SS and VIS). Conversely, L6IT_IPCs showed the strongest FB tendency, aligning with their axonal projections to deep cortical layers and somata predominantly in high-order brain regions (PFC, AI, and MO).

Our WNM cell analysis showed that brain hierarchy is best predicted by target number and TargProb rather than TargStren, with higher-order areas receiving more projections and sensory regions ranking lower. Most cell types exhibited FF or FB biases, notably L4 cells favoring FF and L6IT_IPCs favoring FB, aligning with their projection patterns and soma locations. These findings refine the framework for studying hierarchy at the single-cell level.

### Comparative analyses of the C-C and C-TH connectomes detected by WNM cells and bulk anterograde injections

Bulk anterograde AAV tracing studies have been instrumental in identifying projection targets within neuronal connectomes[35, 77, 78]. However, they face limitations, including false-positive labeling from contamination, inability to distinguish passing fibers and incomplete coverage. Single-cell WNM data overcomes these issues by fully reconstructing dendritic and axonal structures, enabling high-fidelity projectome mapping even with only a few WNM neurons (**Figure 1e**). To refine target identification, we introduced criteria requiring at least one branching node and one ending node, effectively excluding passing fibers while detecting weaker targets (TargStren < 1000 µm) previously overlooked[6, 40]. Additionally, unlike bulk anterograde AAV injections, which often leave ‘empty’ gaps or fail to represent all cell types in a source area, our approach samples diverse neuron types across a source cortical area, ensuring comprehensive representation of both local and long-range projections and dendrites (**Figure 8a–d**).

**Figure 8.**
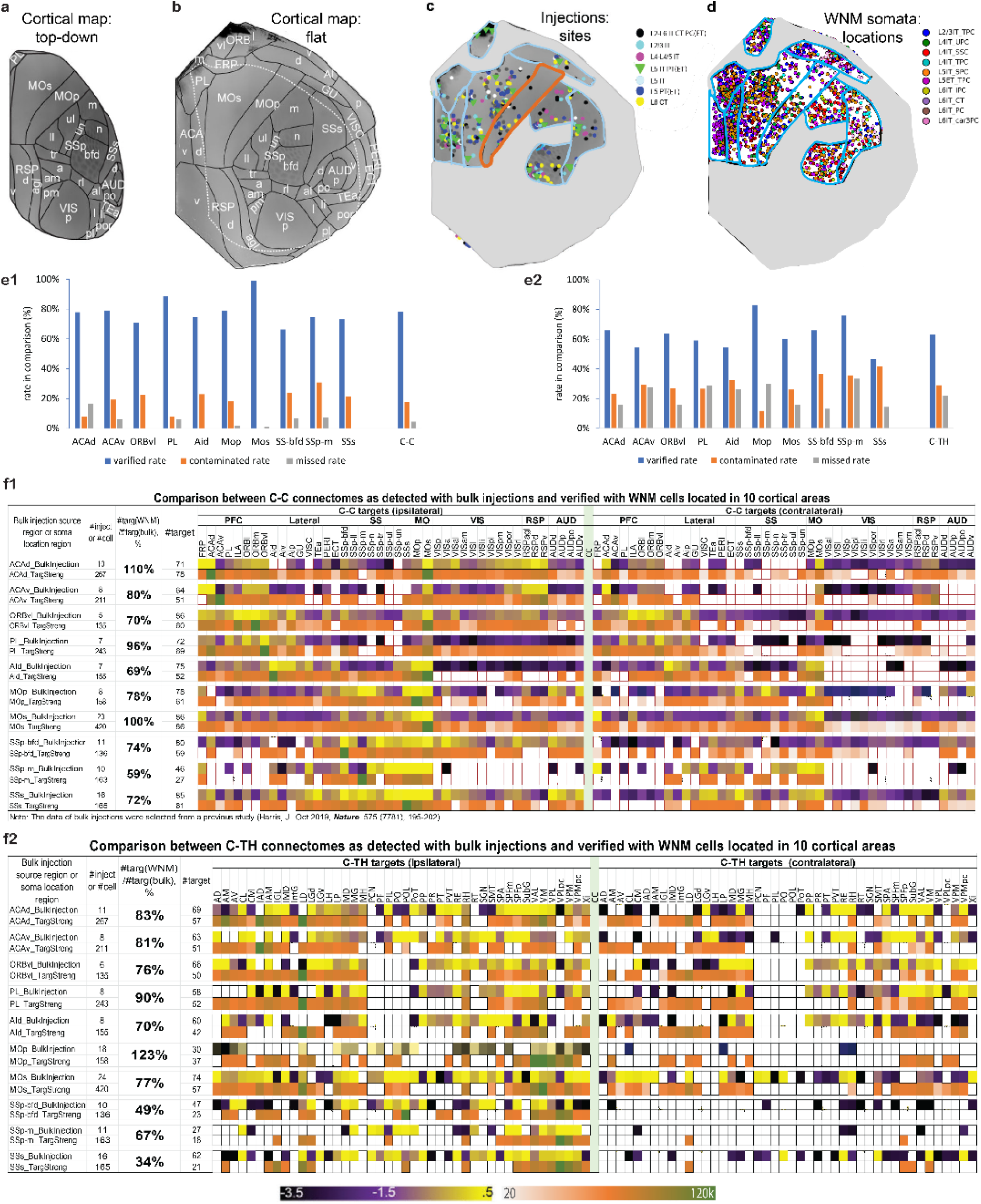
Comparison between projectomes derived from anterograde bulk injections and WNM cells. **a.** Top-down cortical map of the mouse brain. **b.** Cortical flatmap of the mouse brain. **c.** Distribution of anterograde bulk injection sites across ten cortical areas, adapted from our previous study [35]. These cortical areas are outlined on the flatmap. Laminations and cell types of injection sites are defined by transgenic mouse lines color-coded differently (Tg or WT mice). MOp is outlined in orange to highlight its insufficient bulk injection coverage and inadequate representation of labeled cell types. **d.** Flatmap showing locations of WNM somata distributed within the same ten cortical areas shown in (**c**). Ten cell types are color-coded differently. **e.** Validation of C-C (**e1**) and C-TH (**e2**) projection targets. Comparison of three rates across cortical areas: verified rate (number of targets consistent between WNM cells and bulk datasets/total examined targets of 86 for C-C and 88 for C-TH, including zero values), contaminated rate (targets detected only from bulk injections/total bulk injection targets), and missed rate (targets detected only from WNM data/total WNM targets). **f.** One-on-one comparisons of projection targets in the C-C (**f1**) and C-TH (**f2**) connectomes between anterograde bulk injections and WNM cells from ten cortical areas revealed notable discrepancies, shown in two rows for each source area pair: projection volume from bulk injections (**upper**, color-coded by log NPV ranged 10^–3.5 to 10^0.5) and TargStren from WNM cells (**lower**, color-coded by average AxL/C), with zero values outlined in white for both. Particularly in the C-C connectome, some entire contralateral functional regions identified by bulk injections were not all verified by WNM data, suggesting contaminations. Specially in the C-TH connectome, MOp where the target number identified by WNM data was higher than bulk injections, indicats insufficient bulk injections and inadequate cell type labeling representation as shown in (**c**).

To compare C-C and C-TH connectomes derived from extensive bulk anterograde injections, WNM cell targets in each area were matched one-to-one with those from 10 source cortical areas previously published [35] (**Figure 8e, f**). On average, target numbers identified by WNM data were fewer—81% and 75% of target numbers detected by bulk injections for the C-C and C-TH connectomes, respectively (**Figure 8f1, f2**). In some cases, however, WNM cells identified more targets than bulk injections, likely due to insufficient injection coverage or inadequate cell-type labeling in the bulk injection data. For example, in the MOp of the C-TH connectome, WNM cells detected 123% of the bulk injection targets, consistent with such limitations (see **Figure 8c**).

A closer comparison revealed not only contamination (targets reported only by bulk injections) but also missing targets (reported only by WNM cells) across areas (**Figure 8e, f**). Notably, areas where WNM cells identified more targets than bulk injections—such as ACAv and MOp—also had the highest missing rates (17% and 65%, respectively), reinforcing the likelihood of incomplete injection coverage or insufficient labeling of representative cell types. After excluding contamination and missing targets, an average of 77% of total reported (including negative ones) by bulk injections in the C-C connectome and 63% in the C-TH connectome were verified consistent with the reported by WNM cell data.

In the C-C connectome, several notable cases of contamination were identified through direct comparisons between bulk anterograde AAV injection and WNM datasets (**Figure 8e, f; Supplementary Figure 7**). The injection site contamination (encroaching into neighboring cortical areas was most frequently found, especially small cortical areas. For example, WNM cells in AId did not project to the VIS or RSP regions, yet these entire regions appeared as positive targets in bulk injections (**Supplementary Figure 7a1, a2**). This likely resulted from contamination of the claustrum, which lies just beneath AId and directly innervates VIS and RSP [64][79]. Contralateral contamination was also frequently observed. Bulk injections near the midline—such as in ACAv, PL, and ORBvl—may spread across the longitudinal fissure, resulting in contralateral labeling. However, this did not fully account for the contralateral labeling seen in distal regions like AUD, VIS, and RSP following injections in ORBvl and MOp, where the injection sites were not near the midline (**Figure 8f1; Supplementary Figure 7b1, b2**). Retrograde labeling was an unlikely explanation for this, as such cells typically do not show labeling of their own projections [65, 66]. Instead, a more plausible source is the spread of labeling agents via the corpus callosum [67]. For instance, ACAv injections led to contralateral labeling in VIS, and similar patterns were observed in MOp, SSp-bfd, and SSs, likely due to dense callosal pathways between sensory and motor areas (**Figure 8f1**).

Our refined C-C and C-TH connectomes provided more accurate neuronal projectome mapping than bulk injections, which are prone to contamination and incomplete labeling. Single-cell WNM reconstructions eliminated false positive targets, captured weak projections, and better represented diversified cell types, ensuring high-fidelity projectome mapping.

## Discussion

Recent advances in neuroscience have greatly enhanced our understanding of molecularly defined neuronal cell types in the cerebral cortex [1–4], which have been further linked to the whole neuron morphology of major cell categories [6]. However, it still cannot fully capture the regional specificity and morphological diversity of excitatory neuron types. Furthermore, a gap remains between these molecular-WNM classifications and the classical morphological categorization of cortical neurons, traditionally based on brain slice studies using histochemical and Golgi staining techniques, which have provided a foundational understanding of cortical circuitry for over a century [9–30].

To bridge this gap, current study integrates molecular and morphological features to classify ten cortical excitatory cell types. This classification includes seven IT types, L5ET_TPC, L6CT_NPC and L6b_HPC. Through a quantitative analytical approach, this innovative identification of cortical excitatory cell types significantly advances our understanding of organization principles of neuronal circuitries.

### Connectomes formed by single WNM cells across six functional cortical regions

Understanding the intricate networks of C-C and C-subC connectomes is fundamental to neuroscience, as these pathways underpin numerous brain functions. Various methodologies have been used to map these connectomes, including bulk injections [35, 78], electro-physiological and opto-physiological techniques[80, 81], functional MRI [82], and computational modeling [83, 84]. Recent advancements have enabled the reconstruction of individual neurons in specific cortical areas, providing detailed insights into neuronal morphology and connectivity [6, 41, 85–88]. Current study advances the cortical classification by identifying ten morphological types of excitatory neurons across 15 areas spanning six functional cortical regions, coupled with systematic quantitative analyses. This WNM-based classification offers novel insights into targeting patterns, overall targeting power, convergent projections, and topographic and hierarchical organizations.

Our WNM cell analyses reveal that the C-C and C-subC connectomes are organized into four distinct targeting patterns primarily shaped by different groups of cortical excitatory cell types: **Cortical-Dominant Targeting Group:** Comprising L2/3IT_TPCs, L5IT_SPCs, and L6IT_IPCs, these cell types predominantly target cortical areas bilaterally, with stronger projections ipsilaterally. Their projections are highly organized spatially, with L5IT_SPCs exhibiting the most extensive bilateral connectivity, emphasizing their central role in C-C communication.

**Subcortical-Dominant Targeting Group:** Including L5ET_TPCs and L6CT_NPCs, these cells project strongly to subcortical regions, particularly the thalamus, with high specificity and strength. Their targeting patterns highlight the modular and hierarchical organization of information flow between cortical and subcortical areas within the C-subC connectome.

**L4 Cell Group:** Restricted to sensory regions, these cells show primarily localized ipsilateral targeting.

**L6 Cell Group:** L6IT_car3PCs and L6b_HPCs exhibit selective distribution and distinct targeting preferences, with L6b_HPCs’ targeting within ipsilateral hemisphere prominently layer 1.

### The overall balance in targeting power between the C-C and C-subC connectomes

A striking finding of this study is the overall balance in targeting power between the C-C and C-subC connectomes, despite their distinct dominant cell types and projection patterns. When all cell types were pooled, the average targeting probabilities and strengths were nearly equivalent between the C-C and C-subC connectomes, with total target numbers and total TargStren also closely matched. However, when analyzing individual cell types with target numbers and total TargStren, distinct cortical-dominant (C-C) and subcortical-dominant (C-subC) targeting patterns emerged. Notably, when assessing average TargProb and TargStren at the cell type level, most cell types exhibited enhanced targeting power compared to specific non-dominant targets.

As indicated by this finding, a global balance in projectomic power underscores the complementary and balancing roles of C-C and C-subC connectomes in integrating and distributing neural signals. At the same time, distinct targeting patterns at the cell type level and selective enhancement of non-dominant targets maintain specialized processing pathways. These findings highlight the sophisticated organization of cortical excitatory neurons in both C-C and C-subC connectivity, reflecting a balance between functional integration and targeting specificity. Interestingly, studies on human brain function have similarly reported a balance between C-C and C-subC connectomes, emphasizing their integrated contributions to overall brain function[89, 90]. The intricate balance and specificity revealed by single WNM data provide a valuable foundation for future large-scale neuronal network modeling.

### Convergent projections in C-C and C-subC connectomes

Playing a crucial role in brain organization and function, convergent projections remain unexplored at the single-cell level despite extensive studies using anatomical tracing, imaging, electrophysiology, and computational approaches [46–48]. Using high-quality single WNM data, we analyzed convergent projections within C-C and C-subC connectomes, revealing intricate specialization and integration patterns in highly convergent regions: TH, CP, and MOs. Overlap scores quantified extensive axonal clustering from diverse cortical areas in these regions, particularly within higher-order functional areas, while projections from sensory regions showed more localized integration with lower overlap scores.

By analyzing projections from 15 source cortical areas to the three convergent regions, we uncovered distinct targeting patterns reflecting their complementary roles: CP and MOs integrate diverse cortical inputs for higher-order processing, whereas TH serves as a subcortical relay, facilitating cortico-subcortical communication. Notably, the cell type groups clustered within these convergent regions closely matched those identified in the C-C and C-subC connectomes at a whole-brain level, further validating distinct targeting specializations of different cell types and supporting the subjective cell classification as well.

In the C-TH connectome, functional modules have been proposed as specialized pathways for diverse brain functions [49–65]. Our results provide the first single-cell-level evidence supporting this concept, revealing six distinct modular clusters in TH targeted by three cell groups originating from PFC/AI/MOs, MOp & SS, and ACAv/RSP/VIS. Meanwhile, these clusters exhibit broad integration among individual modules, emphasizing the interplay between local specialization and global connectivity. This finding offers valuable insights into the balance between specialization and integration in neural processing within a highly convergent region.

### Topographic organization of single cells in C-C and C-subC connectomes

Topographic organization in the brain has been a major focus of neuroscience for over a century [91, 92]. Early studies by Penfield and Brodmann laid the foundation for mapping functional areas [93, 94]. Advances in neuroanatomy, single-cell transcriptomics, electrophysiology, and imaging have since expanded research from macroscopic cortical maps to single-neuron levels [95]. In this study, large-scale, high-quality 3D reconstructions of WNM cells enable precise mapping of neuronal connectivity.

Our findings demonstrate a strong correlation between the spatial arrangement of neuronal somata in source regions and the distribution of axonal centroids in target regions, revealing distinct patterns for cortical and subcortical projections. In C-C connectomes, axonal centroids generally followed the topographic arrangement of somata, preserving input-output relationships across major cortical regions such as MO, SS, and ACAd, with partial deviations in areas like ORBvl, RSP, and VIS. In contrast, C-subC connectomes exhibited more complex topographic relationships, where axonal centroids often deviated from or reversed the somatic arrangement. These significant different topographic patterns between C-C and C-subC connectomes highlight a fundamental principle of neural organization, reflecting the specialized roles of cortical excitatory neurons in processing and transmitting neural signals. Understanding this principle has broad implications for neural circuit modeling, *in vivo* electro- and opto-physiology, and functional mapping studies. For instance, our study provides precise GPS-like information for *in vivo* study using electrophysiological recordings[96].

### Hierarchical organization in the C-C and C-TH Connectomes

For the first time, quantitative analysis of single-cell WNM data provides a direct evaluation of brain area hierarchy based on how frequently an area is targeted by others. By integrating targeting frequency and axonal cluster size, the hierarchy score serves as a nuanced metric reflecting both anatomical and functional aspects of neuronal networks. This approach is consistent with existing theories of hierarchical brain organization, including functional connectivity and network theory [5, 71, 72] [73, 74].

Hierarchy score curves offer clear insights into the relative positioning of target areas within the C-C and C-TH connectomes, corresponding well with their structural and functional attributes. *High-order regions*, such as prefrontal areas (e.g., ACAd and AI), show high hierarchy scores, reflecting extensive interconnectivity and roles in higher-order cognition. *Intermediate-level regions* like MOs, typically associated with motor functions, exhibit unexpectedly high scores, suggesting involvement in planning and decision-making. In contrast*, sensory areas* such as VIS and SS *rank lower*, with AUD at the bottom—consistent with their low overlap scores and positions within projection-defined module clusters in convergent neuronal hubs. A similar trend appears in the C-TH connectome, where sensory-related targets in modular clusters 5 and 6 rank lower, while areas associated with higher-order functions rank higher. Furthermore, the distribution of hierarchy scores in both connectomes shows significant correlations with the number of targets and TargProb, but not with TargStren. This suggests that the breadth and probability of connections are more indicative of a cortical area’s hierarchical role than the absolute strength of its projections.

Additionally, our findings are consistent with our previous study that modeled cortical connectivity as feedforward (FF) or feedback (FB) to predict hierarchical positions of cortical areas and network modules[5, 35]. Using laminar termination patterns of C-C and C-TH projections, we obtained distribution of the hierarchy differences for each cell type projections and found that most cell types exhibited FF or FB biases in their projections. Notably, L4 cells favor FF and L6IT_IPCs favor FB directions, aligning with their projection patterns and soma locations.

### Newly defined cell types enable revealing the relationship between projection cell types and their gene expression profiles

In order to gain deeper insight into whether a morphological cell type has a distinct molecular signature [5][7], precise identification of morphological types of molecularly defined cells is essential. In this study, morphological classification was performed by selecting cells based on completeness of full structural reconstruction rather than constrained by the cell type specific for a transcriptomic marker. The WNM analysis revealed that dominant expression of a transcriptomic marker corresponds to specific morphological types but does not follow a strict one-to-one relationship. While marker genes used to generate a Cre driver line predominantly label a single type, they may also label multiple other types with significantly lower probabilities. Meanwhile, a single morphological type can express multiple dominant marker genes. In minor cases, a single marker gene may be dominantly expressed in two cell types. For example, the marker gene *Penk* is expressed in both layers 2/3 and 6a of the cerebral cortex [29] (https://www.jax.org/strain/25112) leading to labeling of both populations in the Cre driver line. Our study further clarified that *Penk* predominantly labels two PC types—L2IT_TPC, distinct from L3IT_TPC, and L6IT_IPC, distinguishable from other layer 6 cell types—identifying distinctions that were previously undefined.

### Refining C-C and C-TH connectomes at a single-cell level: insights beyond bulk injections

The comparison between single-cell WNM data and bulk anterograde AAV tracing-derived C-C and C-TH connectomes highlights the limitations of traditional methods while demonstrating the refined insights gained through single-cell reconstructions. WNM reconstructions provide a detailed view of dendritic and axonal structures, serving as a ground-truth representation of projection structures in C-C and C-TH connectomes. Although WNM data often identifies fewer targets than bulk anterograde injections, they are more accurate and reliable, particularly in detecting weak or spatially confined projections that bulk injections may miss. Under the condition of inadequate injection or cell type coverage in bulk anterograde injections, WNM data identifies more targets. Notably, only a few WNM cells of a specific type are needed to reveal the same projections observed in a bulk anterograde injection using a Cre-driver line specifically labeling the same cell type.

WNM reconstructions not only confirm that identified targets are genuinely innervated by originating neurons but also reveal contamination mechanisms inherent in bulk injections. These contaminations often result from projection overlaps in neighboring regions or unexpected labeling spread such as through callosal structures. Insufficient coverage and inadequate labeling of representative cell types further compromising accuracy. High-quality WNM data thus provides a crucial foundation for more precise and detailed explorations of brain connectivity, particularly in guiding *in vivo* electro- and opto-physiological recordings and modeling neuronal networks.

### Summary

By reconstructing complete dendritic and axonal structures of diverse excitatory neuron types across multiple cortical regions, this study provides a comprehensive framework for understanding C-C and C-subC connectomes at the single-cell level, bridging the gap between molecular classifications, whole-neuron morphology (WNM), and classical morphological categorization. We identified and investigated distinct targeting patterns, convergent projectoms, topographic organization, and hierarchical structures in the C-C and C-subC connectomes at single-cell level.

One of the key findings was the balance in targeting power between C-C and C-subC connectomes, despite their distinct cell type compositions and projection patterns. The analysis of convergent projections revealed specified as well as highly integrated functional hubs in CP, TH, and MOs, while six modular TH clusters were identified, reflecting specialized yet integrative roles in C-subC communication. Additionally, the topographic organization of single neurons demonstrated different principles of spatial arrangement governing cortical and subcortical connectivity. Furthermore, based on single-cell targeting properties, our proposed hierarchy score provides a refined metric for evaluating cortical and thalamic hierarchical positioning, and most cell types were verified to exhibit clear FF or FB biases.

By leveraging single-cell WNM reconstructions, this study advances beyond traditional bulk anterograde injection methods, offering higher precision, reduced contamination, capturing weak or spatially confined projections, and improved resolution in mapping brain connectivity. These findings not only refine existing connectoms but also lay the groundwork for future large-scale neuronal network modeling and *in vivo* electro- and opto-physiological studies, ultimately contributing to a deeper understanding of the structural and functional organization of the brain.

## Methods

### Neuron sparse labeling and fMOST imaging

Neuron sparse labeling was achieved using transgenic mouse lines in combination with viral drivers and reporters, as detailed in **Supplementary Table 2**. The expression patterns of many transgenic lines are publicly available in the AIBS Transgenic Characterization database (http://connectivity.brain-map.org/transgenic/search/basic). For transgenic approaches, Cre driver lines were crossed with GFP-expressing reporter lines to achieve sparse yet robust labeling of genetically defined cell populations, as previously described [6, 97]. In CreERT2 lines, optimal tamoxifen doses were determined individually and reported in our earlier work[6]. To further expand the labeling of diverse cell types, viral strategies were employed. The enhancer-based viral driver AiP1046, utilizing mscRE13 (enriched in L6 IT cells), was used to drive Cre expression[98]. Sparse yet strong labeling was achieved using Flp-, Cre-, and Flp/Cre-dual-dependent viral reporters targeting specific recombinase-expressing populations. The Flp-dependent viral reporter (pAAV-TRE-fDIO-GFP-IRES-tTA, Addgene plasmid #118026; http://n2t.net/addgene:118026; RRID:Addgene 118026) was generously provided by Minmin Luo. Cre- and dual-dependent reporters were designed in-house and will be deposited to Addgene. To optimize viral delivery routes and doses, we employed serial two-photon tomography (STPT) for rapid brain-wide screening of sparse and effective labeling. Detailed delivery routes and doses for each case are provided in **Supplementary Table 2**.

fMOST imaging was detailed in our previous study[6]. Briefly, to obtain whole-brain image datasets of sparsely labeled cells using a two-photon fluorescence micro-optical sectioning tomography system (2p-fMOST), a GFP-labeled brain was first embedded in resin. GFP fluorescence was recovered via chemical reactivation by adding Na₂CO₃ to the imaging water bath. A line-scanning block-face imaging system was then used to maximize imaging speed. After each imaging cycle, the top 1 μm of tissue was removed using a diamond knife to expose the next imaging plane. This process generated a 15–20 TB dataset for each mouse brain, comprising ∼10,000 coronal sections at 0.2–0.3 μm xy resolution and 1 μm z-intervals, completed within approximately two weeks.

### Complete neuronal morphology reconstruction and registration to CCFv3

Using 79 mouse brains across 25 transgenic lines, we fully reconstructed 1,419 excitatory neurons, primarily from 15 cortical areas (**Supplementary Table 3**). An unbiased approach was used to reconstruct complete dendritic and axonal structures from any well-labeled cortical cell in sparsely labeled brains with Cre-gene expression. For certain cell types, reconstruction was prioritized based on characteristic morphological features to efficiently increase their sample size, regardless of Cre-gene expression. Reconstruction was performed using Vaa3D, an open-source platform integrating TeraFly for multi-scale visualization and TeraVR for immersive, stereo-based neuron annotation, enhancing both precision and efficiency. Morphological quantification was based on SWC-format reconstructions and followed by expert quality control using in-house tools (Arboreta and NeuroReconManager) to ensure structural completeness. This was performed by checking with original image stacks, including correcting errors, reconstructing missing branches, pruning terminals, and removing misconnected segments. For connectome analysis, we introduced the following new criteria: (1) arborizations with axonal lengths exceeding 1000 µm were required to include at least one terminal node; and (2) arborizations between 20 and 1000 µm needed to have at least one terminal node and one second-order bifurcation. These criteria were designed to exclude passing fibers and artificial tracing spikes and to capture weak but meaningful projections (TargStren < 1000 µm) that were often missed in previous studies [6, 40]. Furthermore, by sampling a broad diversity of neuron types across cortical regions, our approach yielded a more comprehensive and representative projection profile compared to traditional bulk injection methods.

The registration of reconstructed cells to CCFv3, as previously detailed [6], was performed using mBrainAligner to align fMOST images to the CCFv3 template. The process included image downsampling, stripe artifact removal, contour-based landmark matching for affine alignment, intensity normalization, and detection of high-curvature landmarks. Iterative deformation and local alignment were achieved using texture, shape context, and deep-learning features with smooth-thin-plate-spline (STPS). Final results were verified and adjusted when necessary. Once aligned, reconstructed neurons and somata were mapped to CCFv3 space. To correct for deviations caused by individual brain variability, expert manual adjustments were applied post-registration for cells near laminar or area borders, based on their characteristic morphological features.

To complement our dataset, we selected 1,455 additional excitatory neurons with similarly detailed structures from the ML and CEBSIT databases, based on the completeness of dendritic and axonal reconstructions (ML: https://ml-neuronbrowser.janelia.org/; CEBSIT: Digital Brain, digital-brain.cn).

### Data preparation, hierarchical clustering, ANOVA statistical tests and random Forest classifier

Apical dendritic, basal dendritic, and axonal features were extracted from SWC files of 2,874 cells across 15 cortical regions using in-house Arboreta’s neuron_statistics.exe extractFeatures program. To improve clustering accuracy and reduce overfitting, we applied biological insights and feature selection techniques to remove redundant or irrelevant features. A correlation matrix was generated using Seaborn’s heatmap to identify features most strongly associated with cell type, resulting in the selection of 21 optimal features for clustering (**Figure 2a**). To enhance cluster separation and visualization across 15 areas, sub-data frames were created for Layers 1–4, Layer 5, and Layer 6. Each dataset was normalized using min-max scaling (i.e., **min-max normalization**) to ensure consistency across feature ranges. The normalized sub-datasets were then uploaded into [Morpheus](https://software.broadinstitute.org/morpheus/), where hierarchical clustering was performed using the one-minus Pearson metric and average linkage method. Cluster diagrams were generated for each area, with cell types color-coded to clearly illustrate clustering accuracy (**Figure 2a**).

For the random Forest classifier (**Figure 2b**), we selected a subset of nine apical dendritic features optimized to maximize classification accuracy while minimizing overfitting. These features include: *height*, *max distance from soma*, *max pathway length from soma*, *total pathway efficiency*, *average bifurcation-to-bifurcation angle*, *average segment length*, *number of segments*, *number of bifurcations*, and *number of tips*. The ten classified cell types were: L2/3IT_TPC, L4IT_SSC, L4IT_TPC, L4IT_UPC, L5ET_TPC, L5IT_SPC, L6CT_NPC, L6IT_IPC, L6IT_car3PC, and L6b_HPC. To assess statistical significance in feature differences across regions, we used the anova_lm function from the *statsmodels* library, with the ’typ’ parameter set to 2 to accommodate unbalanced data across regions. We performed 100 fitted linear regression models to examine pairwise combinations of features and cell types, compiling the resulting p-values into a summary table (**Figure 2c1**) and presenting the feature data as mean ± SE per cell type (**Figure 2c2**).

We initially trained a random Forest classifier (RFC) model on the full dataset spanning all 15 cortical areas, using the selected morphological features as predictors (X) and cell types as labels (y), encoded via a label encoder. The dataset was split into 80% training and 20% testing sets. The trained RFC model was then used to predict cell types on the test set. To assess area-specific performance, we trained 15 independent RFC models using area-specific data frames. For each area, we calculated the classification accuracy by comparing the predicted cell types against actual labels in the test set. Overfitting was evaluated by comparing training and validation scores, and to mitigate overfitting, we fine-tuned hyperparameters using both GridSearchCV and RandomizedSearchCV from the *sklearn* library. Accuracy scores were computed as the ratio of correctly predicted cell types to the total number of cells in each region (**Figure 2b**).

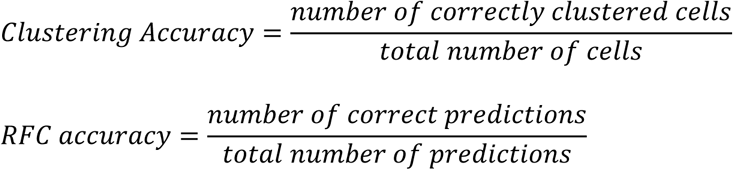

### Quantitative analysis for C-C, C-subC and C-TH connectomes

Axon cluster overlap score

Since neuron reconstruction data consist of a collection of nodes, we treated each axon cluster as a distinct *class*. To evaluate the extent of overlap between any two axon clusters within a target area, we adopted the concept of **class separability** in feature selection as described by Orihuela-Espina et al.[99]. The class separability *S* was given by:

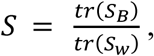

where *S*_*B*_ is the between-class scatter matrix:

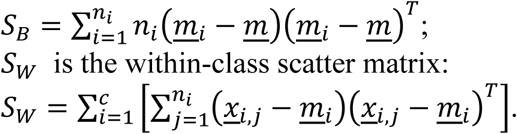

*c*is the number of classes (*c*= 2 in our case), *n*_*i*_is the number of nodes in the *i*-th axon cluster, <u>*x*</u>_*i*,*j*_ is the *j*-th node in the *i*-th axon cluster, *m*_*i*_ is the mean vector of the nodes in the *i*-th axon cluster, *m*is the mean vector of all the nodes in the 2 axon clusters to be evaluated. Our overlap measure *O* is defined as: *O* = *max*(0,1 − *S*), with *O* = 0 indicating no overlap and 1 indicating full overlap. To ensure the accuracy of overlap score, we interpolated our raw data so that every Cartesian grid in the straight path from one node to its subsequent child node was included for the calculation of between-class and within-class scatter matrices.

All formulas as above can be integrated into one formula for the overlap score calculation:

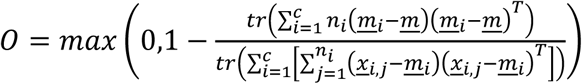

Axon cloud presentation and axon volume estimation

Given the hierarchical nature of neuron reconstruction data (nodes forming line segments in 3D space), we applied kernel density estimation (Gaussian kernel, bandwidth = 0.3) to generate smoothed 3D density distributions for axon clusters formed in TH, CP and MOs by individual neurons from 15 cortical areas. These distributions were used to estimate the volume occupied by each axon cluster and to calculate the average axon density, defined as the total axon length per estimated volume (μm/μm³). The estimated volume was computed as the sum of voxels with density ≥ 1%, an empirically chosen threshold that preserves axonal structural detail without exceeding the target region’s boundaries.

### Topography **ρ**

We evaluated the topological relationship between the locations of the somas from the source regions and their corresponding arborizations in the target regions by calculating the Spearman’s rank correlation coefficient ρ as follows:

Let *S* be the set of soma coordinates from the source region and *C* be the set of the centroids of arborizations in the target region. Note that in order to ensure the centroids of the arborizations are accurate, we interpolated the nodes in every segment so that the node density is uniformly distributed throughout the whole structure. Subsequently, we identified the first principal axes of *S* and *C* by Hotelling transform as 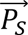 and 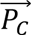 , respectively. Since *S* and *C* demonstrate the largest variance in 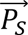 and 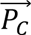, which translates to the widest spatial span of the data set, 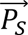 and 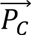, shall be the optimal directions to examine the topological relationship between *S* and *C.* To do this, we obtained the projected sets of *S* and *C* onto 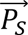 and 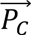 denoted as *S*^′^ and *C*^′^, and then calculated the ρ value of *S*^′^ and *C*^′^. ρ value is ranged from -1 to 1. A negative ρ indicates that somas and arborizations tend to spatially align in the opposite way.

### Hierarchy score calculations

We hypothesized that hierarchical organization within the cortex could be directly reflected in the targeting properties of the C-C and C-TH connectomes. Specifically, if a brain region occupies a higher position in the hierarchy of information flow, it should be more frequently targeted by regions of lower order. Thus, the targeting probability from region *i* to region *j* (*P₍ᵢⱼ*₎), which represents how often region *j* receives input from region *i*, can be interpreted as *i*’s contribution to *j*’s hierarchical status. Our observations showed that larger cortical regions, such as ACAd and MOs, tended to receive more projections than smaller regions. To avoid size-related bias in evaluating hierarchy, we normalized targeting probability using the average targeting length 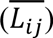, which reflects the extent of the axonal cluster from region *i* to *j*, rather than the physical size of the entire target region. This adjustment accounts for the spatial aggregation of targeting clusters from different sources. Accordingly, the hierarchy score of region *j* was defined as:

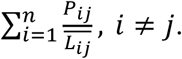

where *n* is the number of cortical regions included in the analysis.

### Classifying the FF/FB-ness of projections from cell types

We used a hierarchy of brain areas, constructed based on laminar source and termination patterns of corticocortical, thalamocortical, and corticothalamic connections, to assess the FF/FB characteristics of cell-type-specific projections [35]. For each connection from area *j* to area *i*, the hierarchy difference was calculated as:

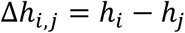

ℎ_*i*_ and ℎ_*j*_ are the corresponding hierarchy levels of areas *i* and *j*, respectively (**Figure 7c**). By grouping connections for each cell type, we generated distributions of hierarchy differences. To evaluate these distributions, we used the Wilcoxon test to compare each one with a normal distribution centered at zero with the same standard deviation. This allowed us to assess whether the observed hierarchy differences deviated significantly from the null expectation. To correct for multiple comparisons, we applied the Benjamini-Hochberg procedure to control the false discovery rate. The significance level of hierarchy difference distributions deviated from the normal distribution was defined at the 95% confidence level. The direction of the lean was defined by the mean of each distribution, while the strength of deviation was reflected in the corresponding *p*-values.

## Supporting information

Wang_bioRxiv-SFigs&STables

## Acknowledgements

We thank Karel Svoboda, Josh Siegle, and Jian-Fan Chen for their valuable comments, suggestions, and insightful questions during the review of our manuscript. We are grateful to Christof Koch for his continued support and sustained interest in this work. We also thank Julie Harris for facilitating access to the bulk injection datasets, and Jayaram Chandrashekar for providing the original ML reconstruction dataset. We thank the Animal Care, Transgenic Colony Management and Lab Animal Services teams for mouse husbandry and tissue preparation. We thank all the members of the Neurosurgery and Behavior team for the viral injection, including those not listed as authors: N. Berbesque, N. Bowles, S. Cross, M. Edwards, S. Lambert, W. Liu, K. Mace, N. Mastan, C. Nayan, B. Rogers, J. Swapp, C. White, and N. Wong. We also thank H. Gu for cloning of the synaptophysin-EGFP viral vector, E. Lee, F. Griffin, and T. Nguyen for intrinsic signal imaging, and J. Royall. This work was supported by the Allen Institute for Brain Science and, in part, by National Institutes of Health grant 5U19MH114830-03 to H.Z. During the early stages of the project, the WMU team also received support from the Blue Brain Project, Switzerland. We thank the Allen Institute founder, Paul G. Allen, for his vision, encouragement, and support.

## Author contributions

Conceptualization: Y.W., H.C.K., S.Y., J.Z., H.Z.; Writing, analysis & figure making: Y.W., H.C.K., X.K., S.Y, P.L., K.Z.; Review & editing: Y.W., H.C.K., X.K., S.Y.,Y.B., S.D., Q.W., J.Z., *S.S.*; Neuron reconstruction: Y.W., X.K., P.L., Y.L., L.E.H., N.C., K.Z., W.Z., C.C., K.C., Z.H., Z.Y., W.X., L.A.; Quality control: Y.W., X.K., P.L., Y.L.; Registration: Y.W., H.C.K., X.K., L.NG., S.W.B., A.L.; Image acquisition: A.L., H.G., Q.L.; Soma marking: Y.W., X.K., Y.L., L.E.H., N.C., R.D., G.W.; Software development: H.C.K, Y.W., L.E.H, N.C, T.K., C.F., B.S., O.G., U.S.; Supervision work: Y.W., H.C.K., P.L., L.NG., K.J., Z.Y.; Requested special analysis: K.Z., E.M.L., M.M., H.C., S.M.; Funding Acquisition: H.Z.; Project Administration: X.K., L.E.,S.S., L.K.,S.S..

