## Supplementary material for "Integrated Classification of Cortical Cells and Quantitative Projectomic Mapping Unveil Organizational Principles of Brain-Wide Connectomes at Single Cell Level": Wang_bioRxiv-SFigs&STables

### Supplementary Figures and Tables

#### Supplementary Table 1

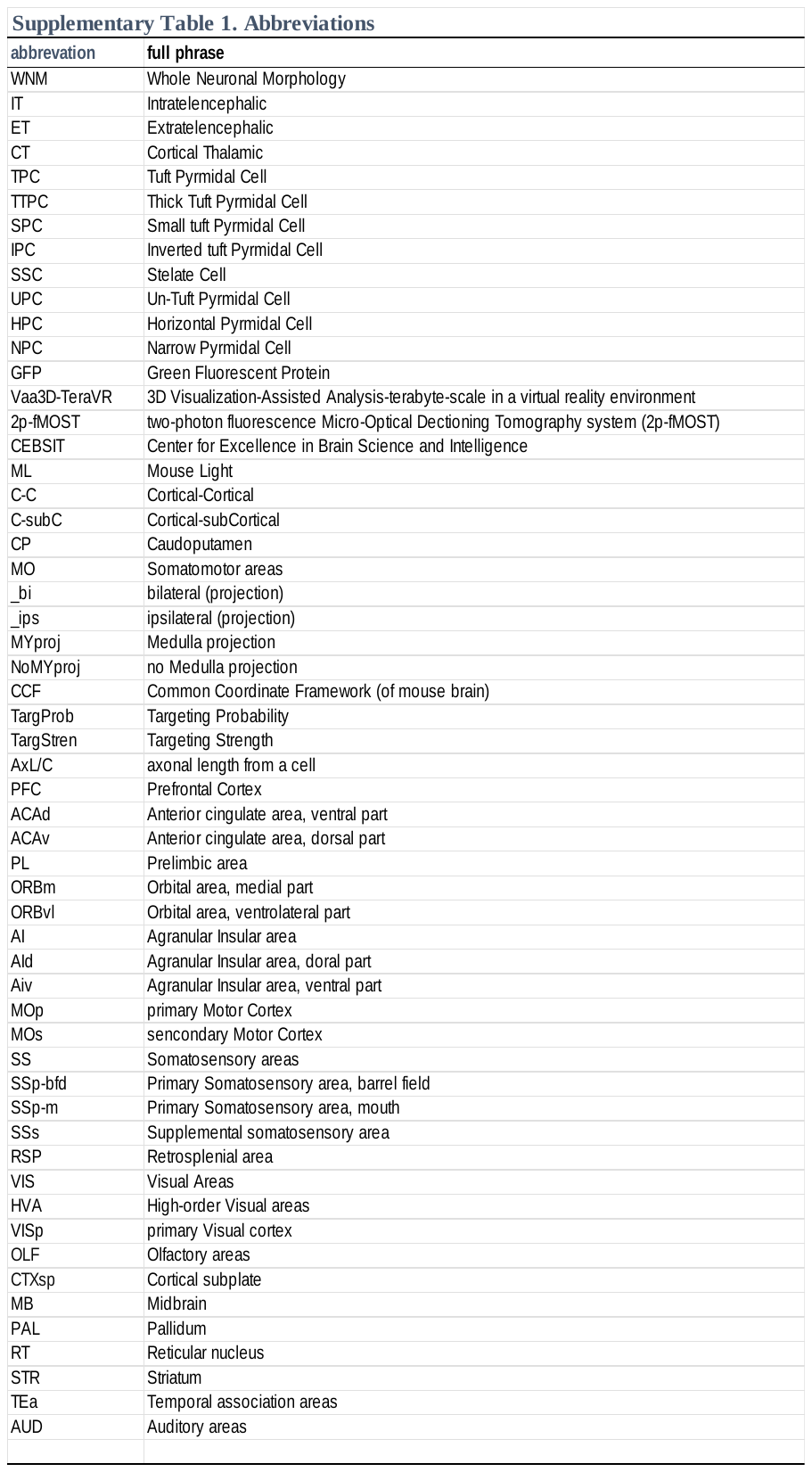

#### Supplementary Table 2

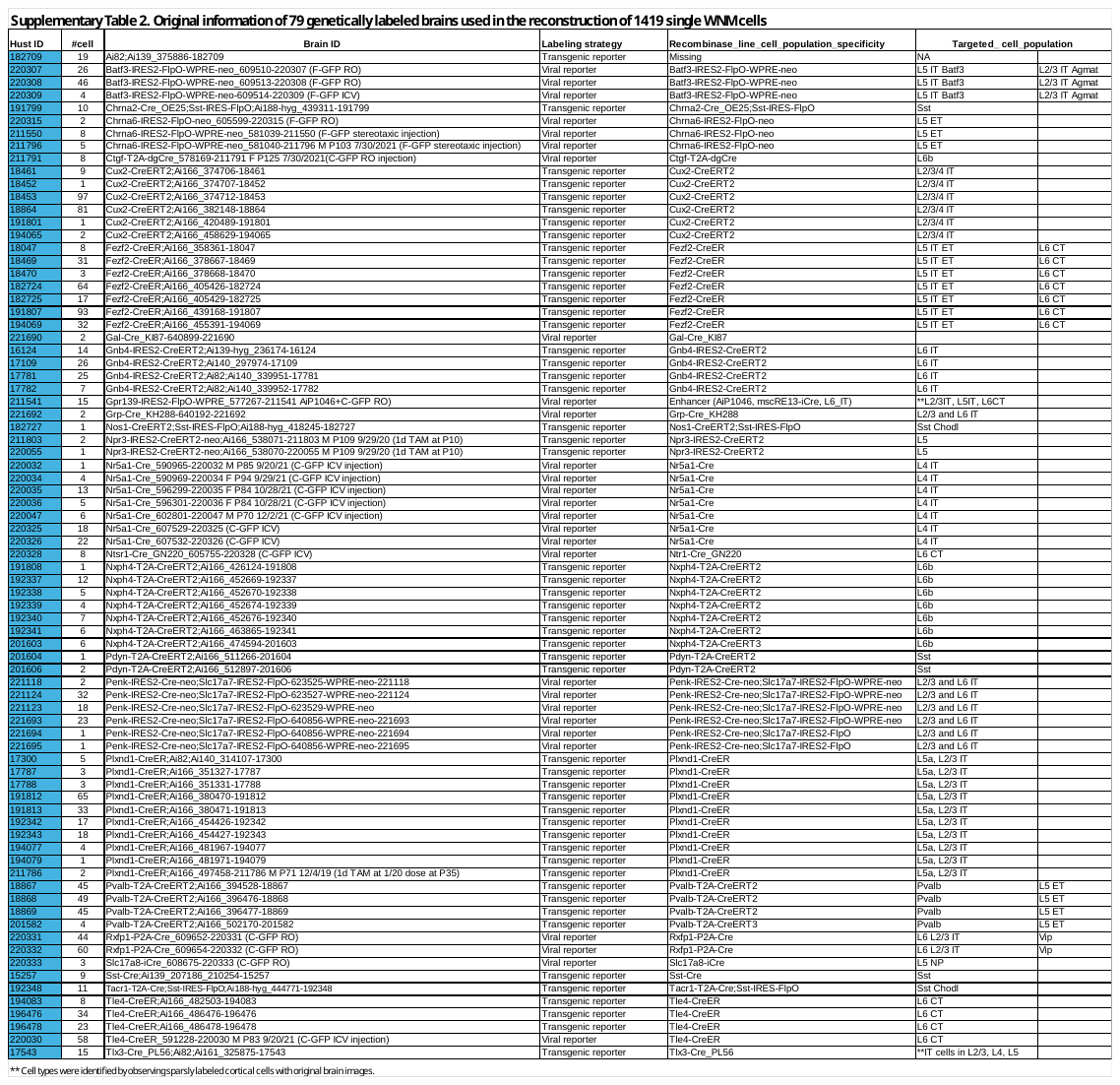

#### Supplementary Table 3 (full information of all cells, will be available when published in a formal journal next)

#### Supplementary Table 4a

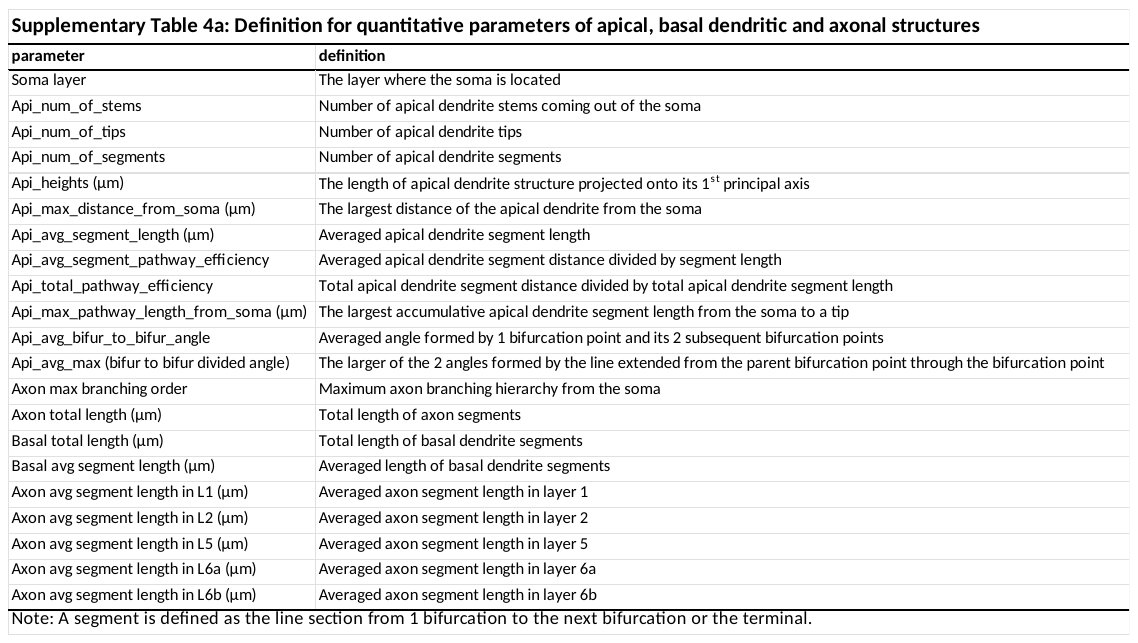

#### Supplementary Table 4b

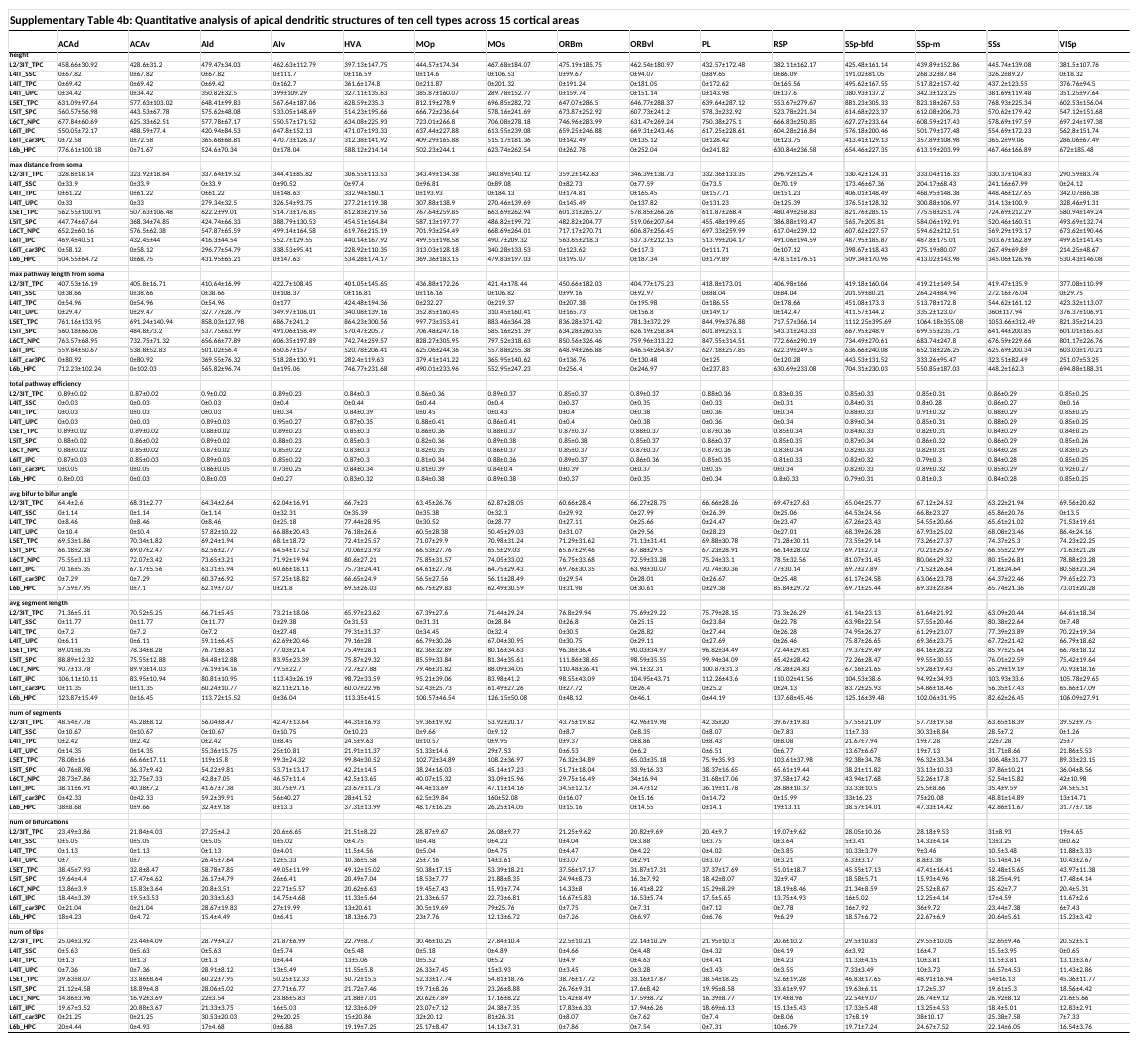

#### Supplementary Table 5

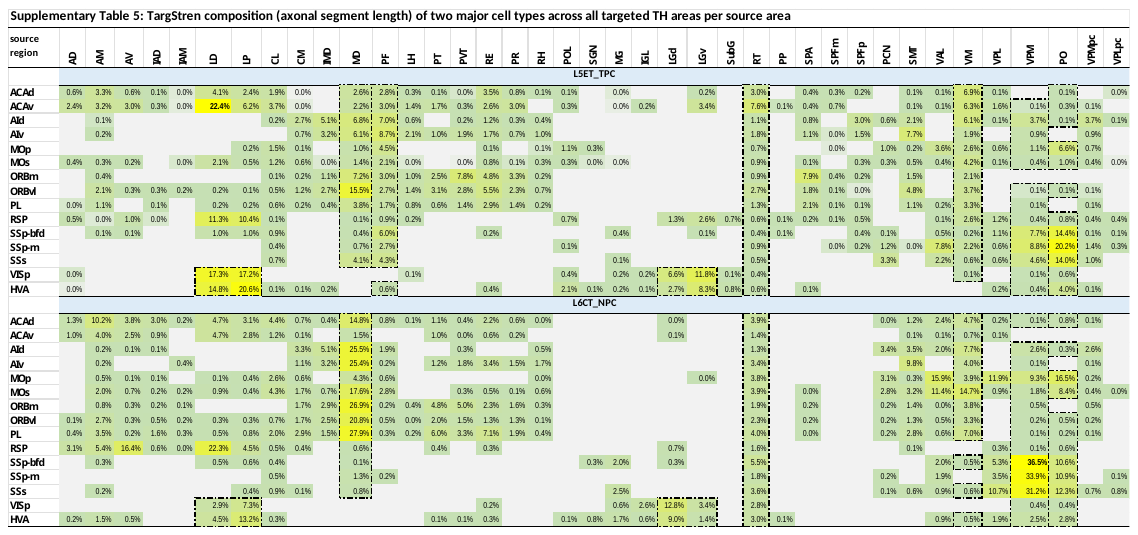

#### Supplementary Figure 1.

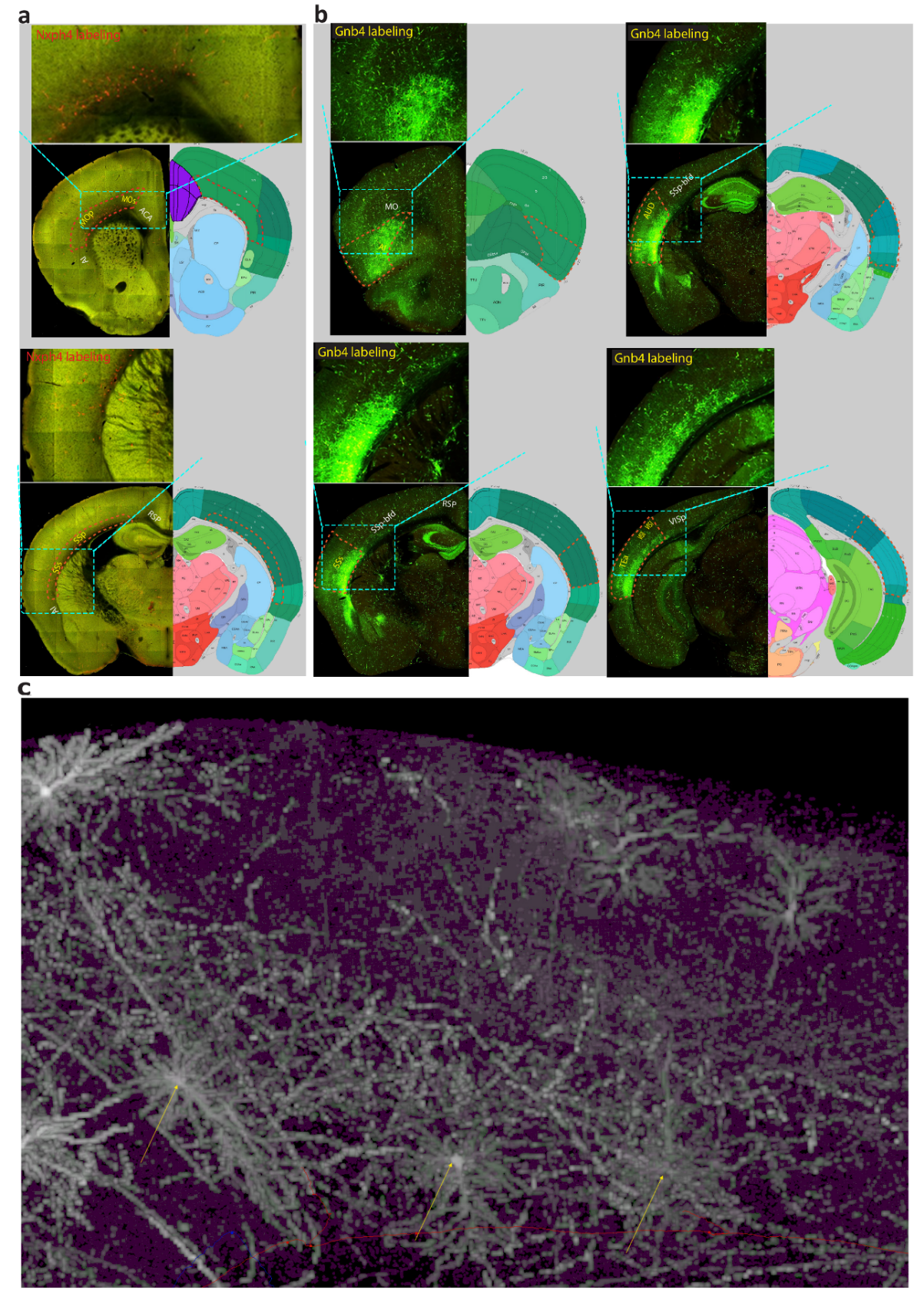

**Supplementary Figure 1**. **Evidence of transcriptomic markers** **for selectively labeled cell types**

**a. Upper panel**: Nxph4 expression was predominantly observed in the lower part of layer 6 (mainly L6b, <https://lims2.corp.alleninstitute.org/drawing_tool?image_series=719201424>) in the MO, but not in adjacent areas such as ACA and AI (positive labeling areas indicated in yellow font within dashed circles; negative areas in white font). **Lower panel**: Similar expression was found in the SS region but absent in neighboring RSP and AI areas.

**b.** Gnb4 expression was observed in layer 6 of the AI, SSs, AUD, TEa, VISl, and VISli regions, but not in adjacent areas such as ACA, MO, RSP, SSp-bfd, and VISp (<https://lims2.corp.alleninstitute.org/drawing_tool?image_series=572389684>).

**c.** Three L6IT_IPC neurons (brain ID: Penk-IRES2-Cre-neo; Slc17a7-IRES2-FlpO-640856-WPRE-neo-221694) were identified by their inverted apical dendrites or an inverted with a big upward dendrite forming bipolar-shaped dendritic clusters, with axonal arbors confined mainly to deep layers.

Note: data for **a** & **b** from <https://lims2.corp.alleninstitute.org>.

#### Supplementary Figure 2

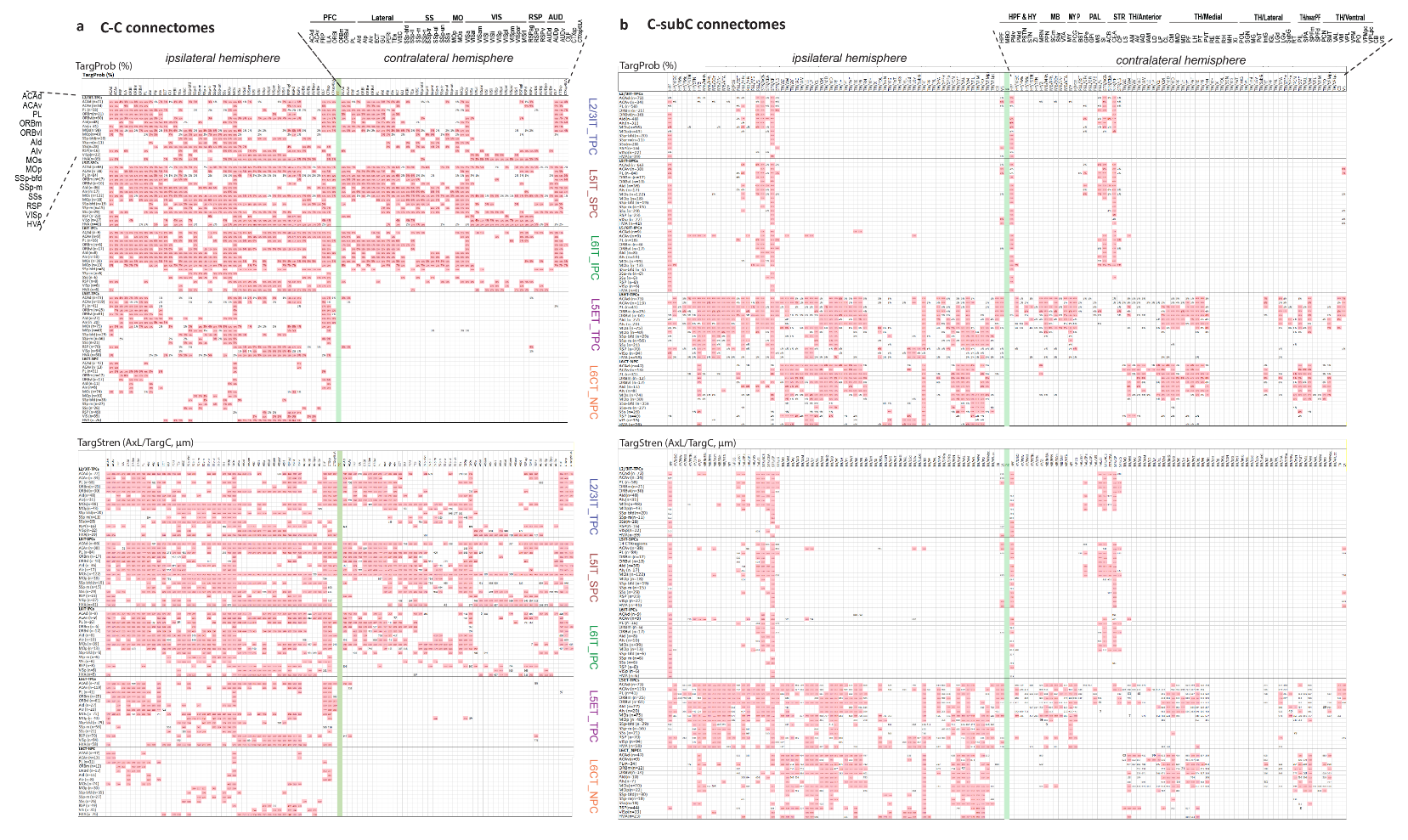

**Supplementary Figure 2.** **Correlation between TargProb and TargStren of weak targets in C-C and C-subC connectomes**.

**a**. Targeting pattens are matched between TargProb and Targstren plots in the C-C connectome: 93% of the total 2,267 targets are highlighted in pink, based on a TargStren threshold of 1000 µm (**lower panel**) and a corresponding TargProb threshold of 2.4% (**upper panel**).

**b.** Targeting pattens are matched between TargProb and Targstren plots in the C-subC connectome, 69% of the total 1,749 targets are highlighted in pink, determined by the TargStren threshold (1000 µm, **lower panel**) and a corresponding TargProb threshold of 6% (**upper panel**).

Note: Targets below the thresholds are displayed in their original values without highlighting, representing weak targets (7% of the total in C-C connectome and 31% in C-subC connectome).

#### Supplementary Figure 3

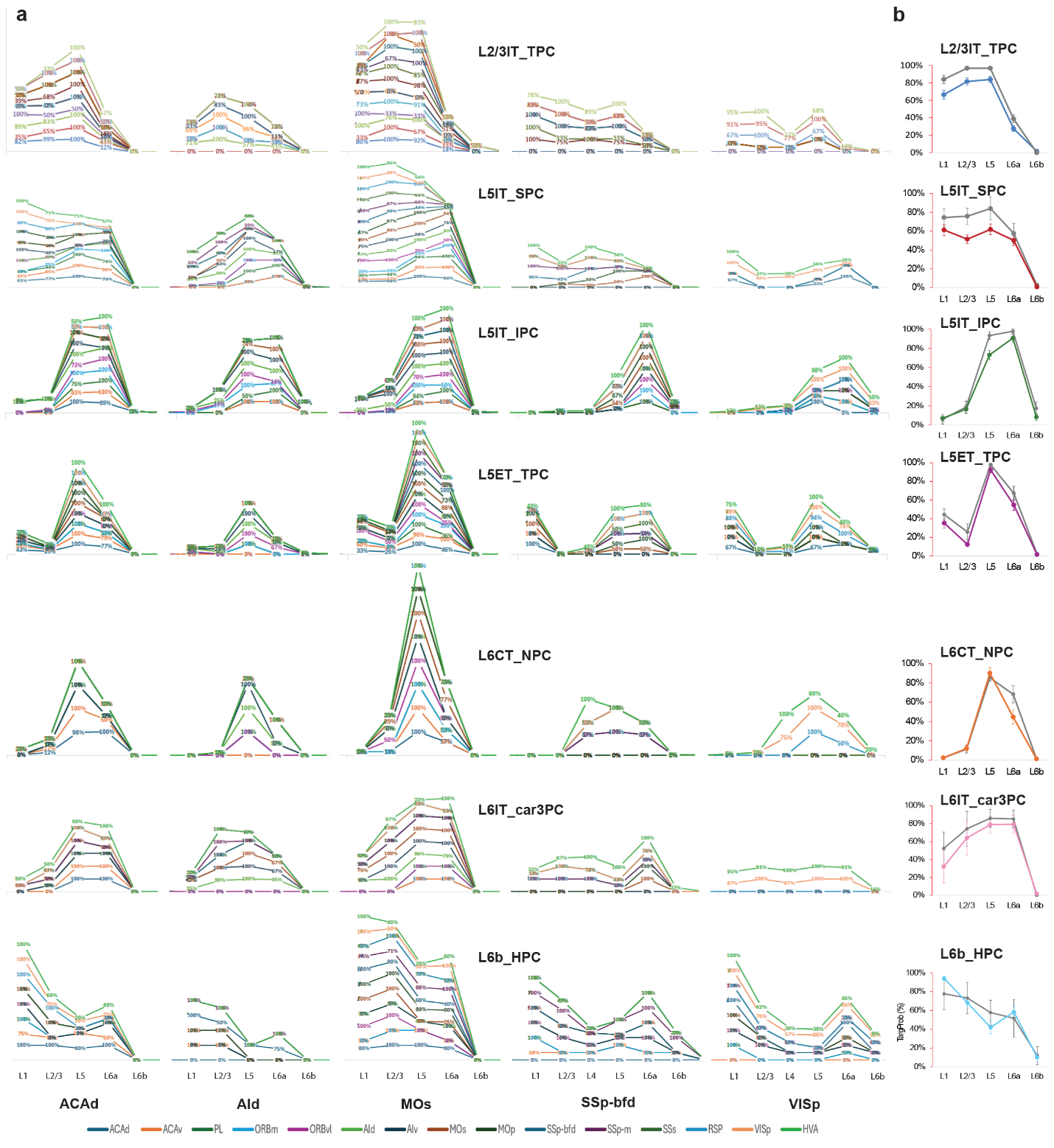

**Supplementary Figure 3. Axonal laminar patterns of seven cell types across 15 source areas targeting five cortical areas.**

**a.** Individual axonal laminar patterns of seven cell types from different source areas to five representative target areas: ACAd, AId, MOs, SSp-bfd and VISp. Specifically, L2/3 IT-TPCs primarily target layers 2/3, 5, and 1, with secondary projections to layer 6a; L5 IT-SPCs predominantly target layers 1, 5, and 2/3, with secondary projections to layer 6a; L6 IT-IPCs mainly target layers 6a and 5, with secondary projections to layer 2/3, and rarely to layer 1; L5 ET-TPCs chiefly target layer 5, with secondary projections to layers 6a, 2/3, and 1; L6 CT-NPCs also mainly target layer 5, with secondary projections to layers 6a and 2/3, and rarely to layer 1; L6IT_car3PCs primarily target layers 6a, 5, and 2/3, with secondary projections to layer 1; L6b_HPCs predominantly target layers 1 and 2/3, with secondary projections to layers 5, 6a, and 6b. Notably, layer 6b is seldom targeted in most areas, except by L6IT_IPC and L6b_HPC.

**b**. Distinctive laminar patterns of the seven cell types are shown with the average TargProb values across the five source areas (grey) and target areas (in the same color coding for target areas as in **Figure 3c2 & Figure 4b3**).

Note: Data are normalized to the maximum TargProb across layers 1 to 6b for each projection in the ipsilateral hemisphere.

#### Supplementary Figure 4

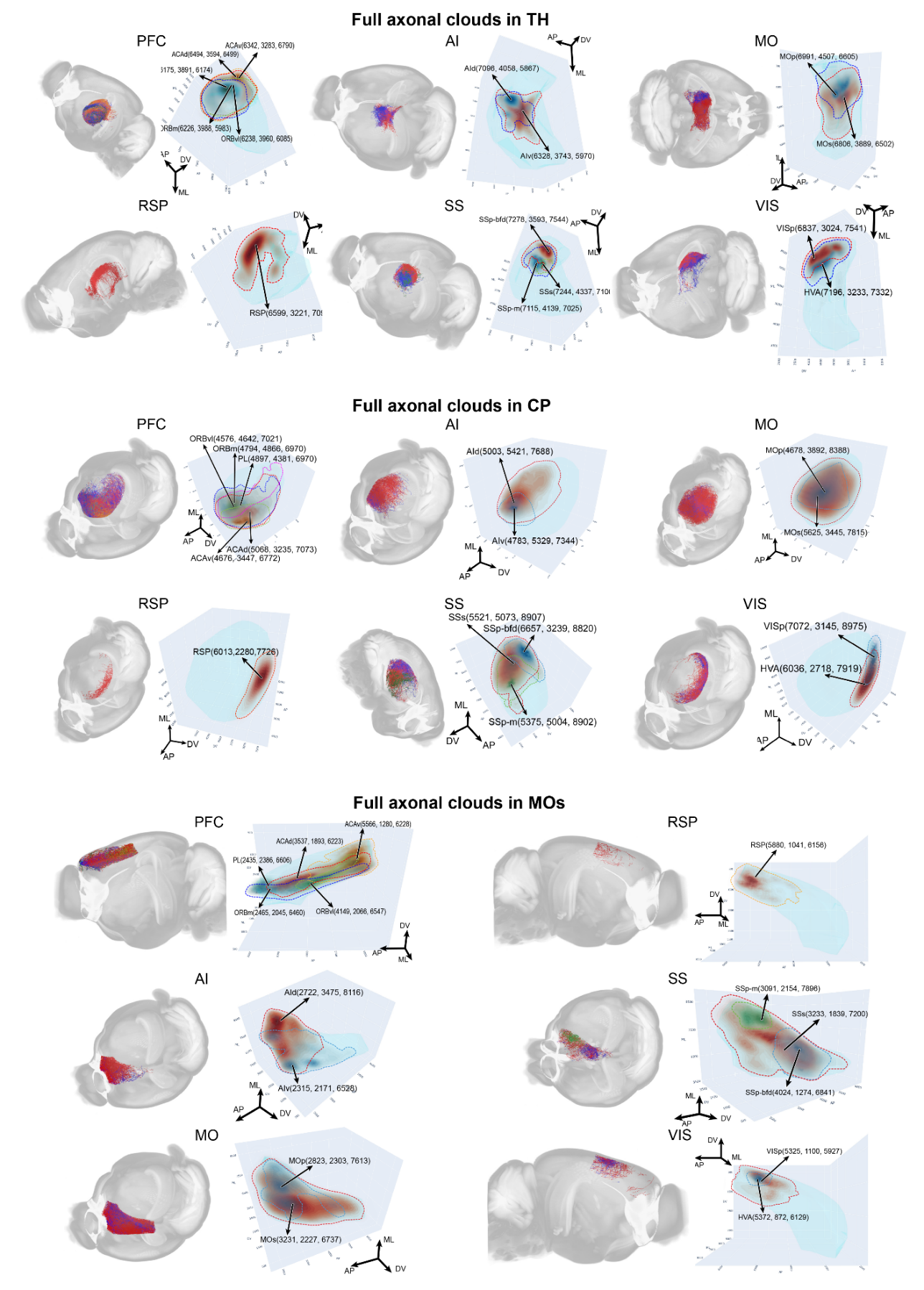

**Supplementary Figure 4.** **Full axonal projection clouds converging in TH, CP, and MOs from cells in 15 cortical areas across six functional regions.** Data is presented by functional region. Convergent axonal segment clouds from different source areas within the same functional region are color-coded and visualized in the CCFv3 template (**Left panel**). In each convergent region, full axonal clouds are outlined in colors matching their respective axonal segment clouds’ colors shown in left panel. The centroid of each axonal cloud is marked by an arrow, with the source area name and XYZ coordinates near the arrowhead (**Right panel**). Notably, axonal clouds from MOs span nearly the entire volume of all three convergent regions. Several other clouds also show substantial expansion, including those from ACAd, PL, AId, and MOp to CP, and from AIv and SSs to MOs.

#### Supplementary Figure 5

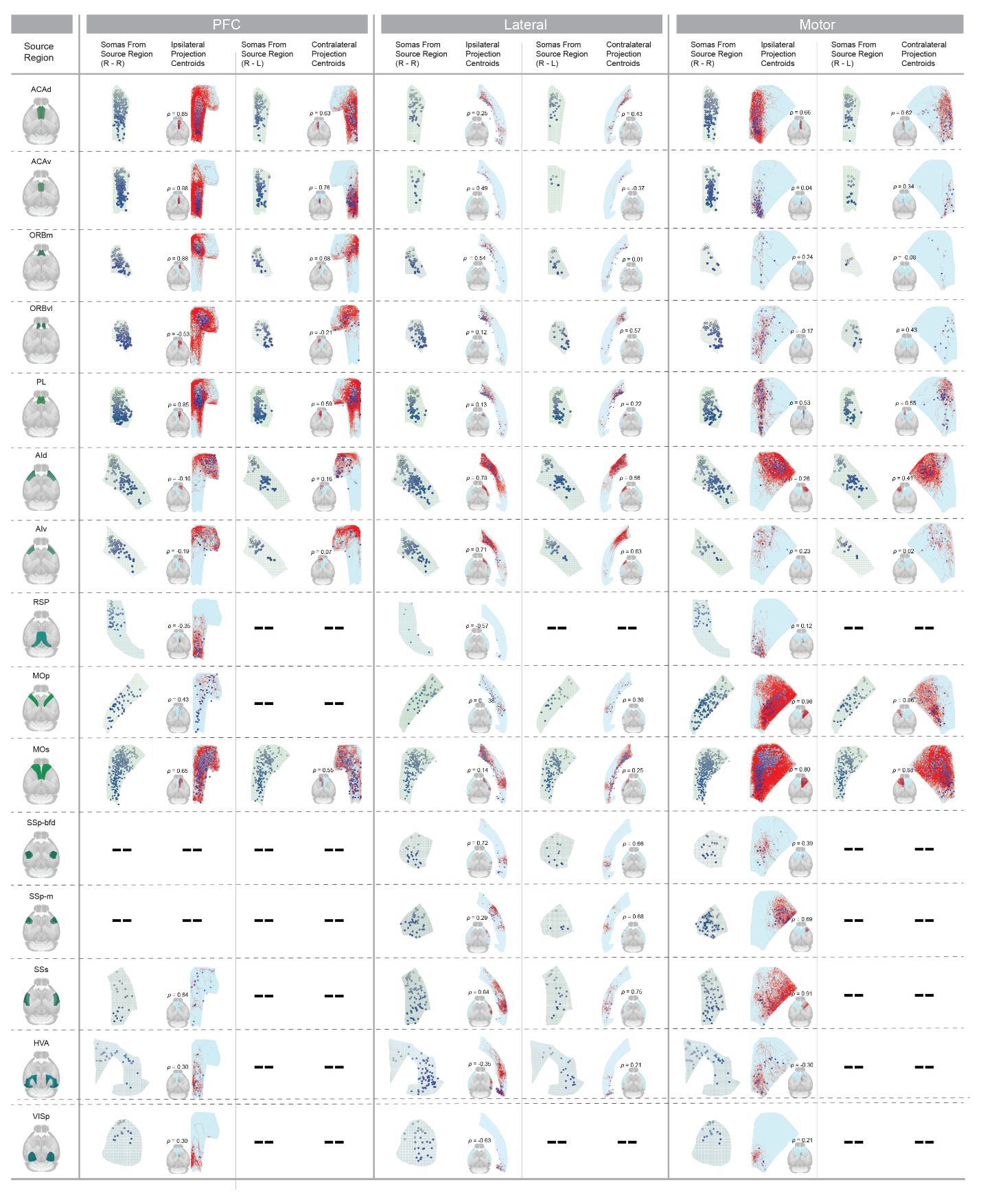

**Supplementary Figure 5. Topography organization in the C-C connectome,** represented by the correlation between soma arrangement of different cell types across 15 cortical areas and corresponding projection centroids of single cells in ipsi- and contra-lateral target regions (PFC, lateral cortex, and MO). Gradient color density is used for visualizing and matching the spatial arrangement of somata in a source region with their corresponding projection centroids in a target region. Projection centroids are overlaid on axonal segments (shown in red), illustrating spatial correlation, location, and the expansion of projecting axonal clouds from source to target regions.

#### Supplementary Figure 6

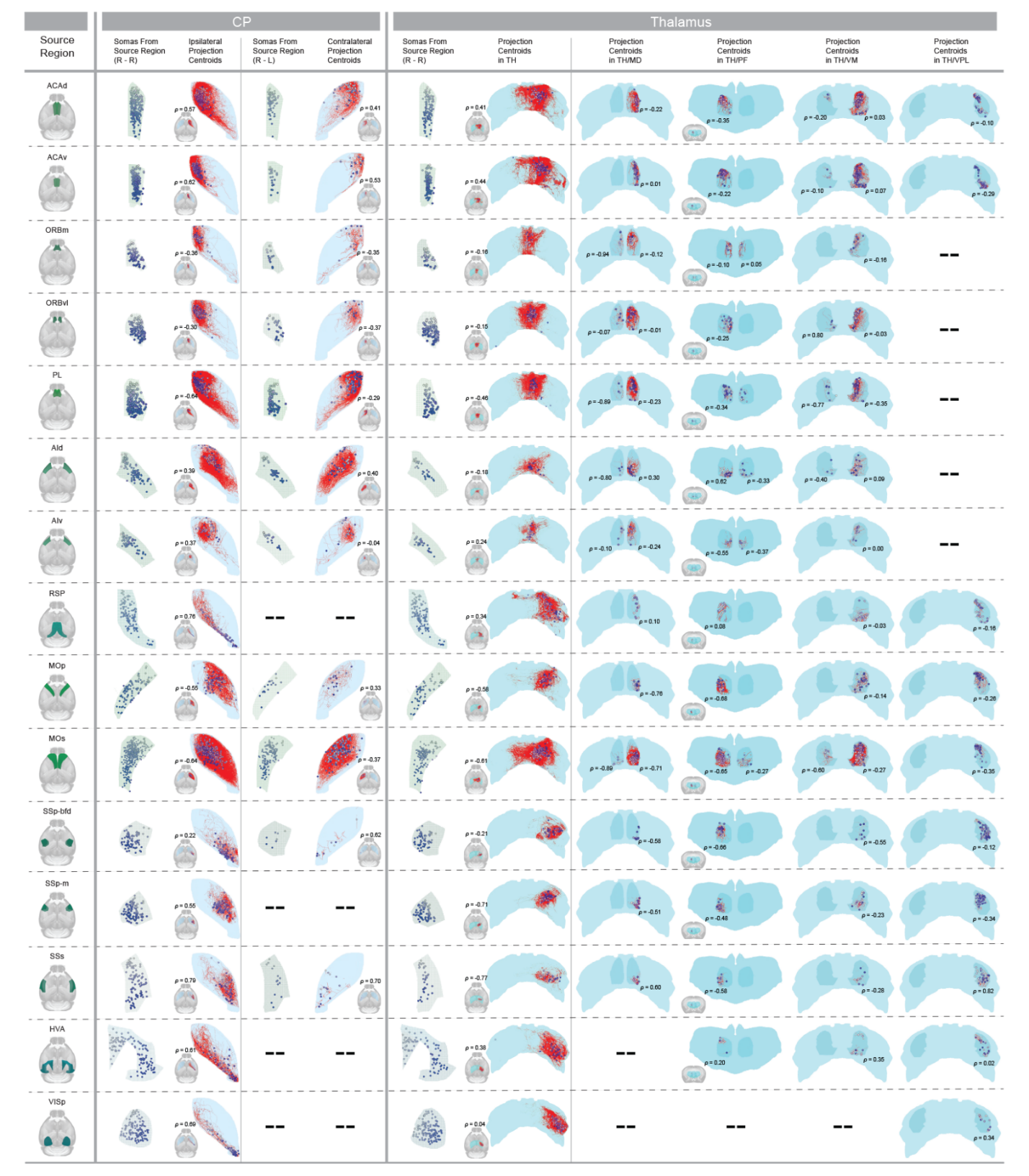

**Supplementary Figure 6.** **Topography organization in the C-subC connectome,** represented by the correlation between soma arrangements of different cell types across 15 cortical areas and corresponding projection centroids of single cells in ipsi- and contra-lateral target regions (CP, TH, and its subareas MD, PF, VM, VPL). Gradient color density is used for visualizing and matching **t**he spatial arrangement of somata in a source region with their corresponding projection centroids in a target region. Projection centroids are overlaid on axonal segments (shown in red), illustrating spatial correlation, location, and the expansion of projecting axonal clouds from source to target regions.

#### Supplementary Figure 7

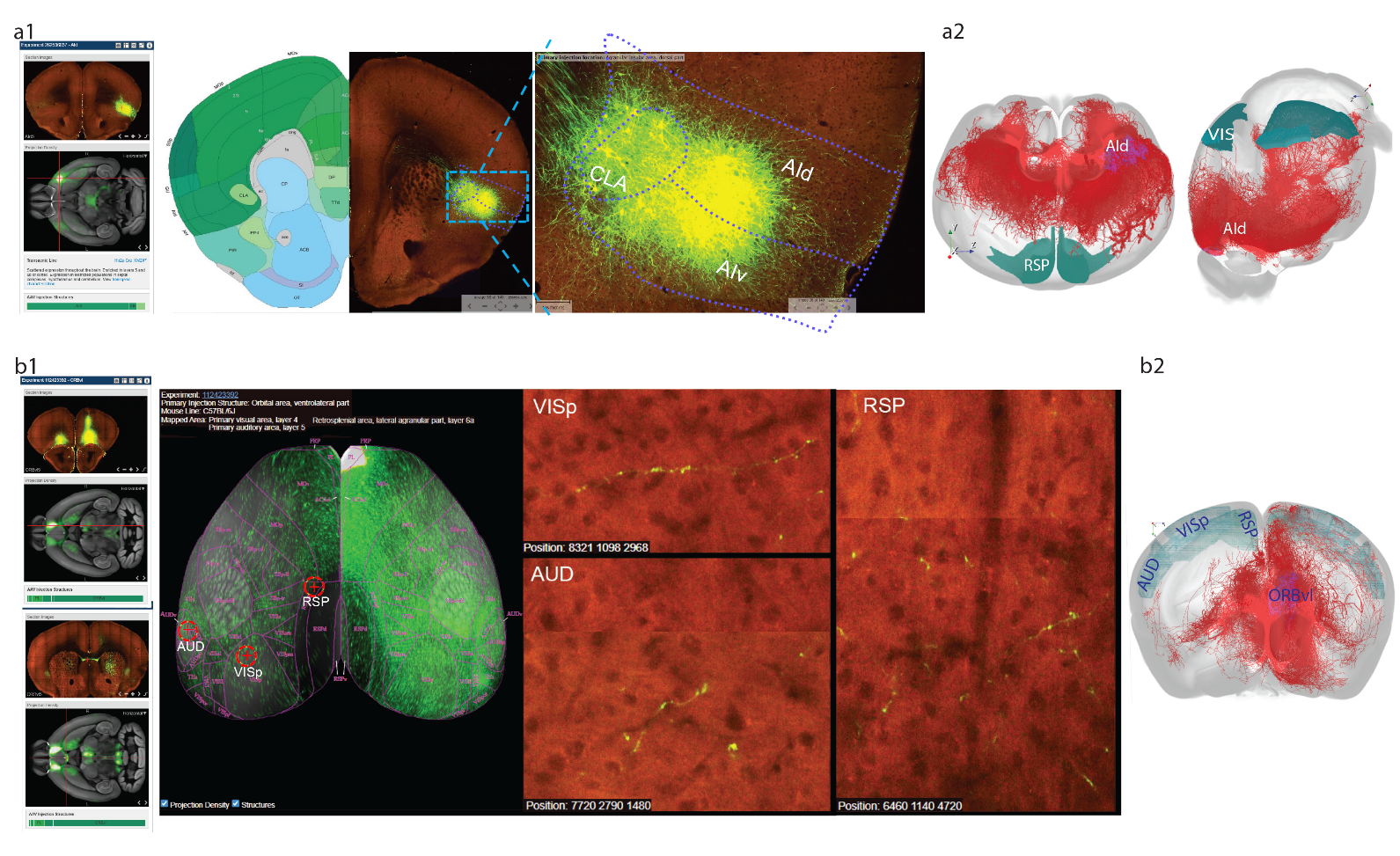

**Supplementary Figure 7**. WNM validation identifies bulk injection target contaminations from adjacent and contralateral areas.

**a**. Validation of contaminations induced by labeling adjacent areas. **a1**. The labeling of a bulk injection in AId (**Left**) contaminated CLA, which sends projections to VIS and RSP (**Middle & Right**, Experiment 262536037 – AId). **a2**. WNM AId cells of different types (n=147) do not project to VIS & RSP.

**b**. Validation of contaminations induced by labeling distal areas in the contralateral hemisphere. **b1**. A bulk injection in ORBvl of the right hemisphere and contaminated contralateral ORBvl (**Left upper panel**), strongly labeled ccg (**Left lower panel**); Contaminated contralateral VISp, AUD, and RSP: red circled crosses mark the three sites on the cortical CCFv3 (**Middle panel**), each accompanied by a zoom-in view showing labeled axonal segments (**Right panel**. Experiment 112423392 – ORBvl). **b2**. WNM ORBvl cells of different types (n=145) have no projections to contralateral VIS, RSP and AUD.

**Note**: All bulk injection data are available from public source (https://connectivity.brain-map.org/).
